# Chromatin state transition underlies the temporal changes in gene expression during cardiomyocyte maturation

**DOI:** 10.1101/2021.12.28.474318

**Authors:** Chia-Yeh Lin, Yao-Ming Chang, Hsin-Yi Tseng, Yen-Ling Shih, Hsiao-Hui Yeh, You-Rou Liao, Chia-Ling Hsu, Chien-Chang Chen, Yu-Ting Yan, Cheng-Fu Kao

## Abstract

Congenital heart disease (CHD) is often rooted in gene expression anomalies that occur during heart development. As cells commit to a specific lineage, chromatin dynamics and developmental plasticity generally become more limited. However, it remains unclear how differentiated cardiomyocytes (CMs) undergo morphological and functional adaptations to the postnatal environment during their maturation. In this work, we sought to identify the regulatory mechanisms controlling postnatal cardiac gene networks. A time-series transcriptomic analysis of postnatal hearts revealed an integrated, time-ordered transcriptional network that regulates CM maturation. Remarkably, depletion of histone H2B ubiquitin ligase RNF20 after formation of the four-chamber heart disrupted these highly coordinated gene networks. Its ablation also caused early-onset cardiomyopathy, a phenotype reminiscent of CHD. Furthermore, we found that dynamic RNF20-mediated modulation of chromatin accessibility during CM maturation was necessary for the operative binding of cardiac transcription factors known to drive transcriptional gene networks. Together, our results reveal how epigenetic-mediated chromatin state transitions modulate time-ordered gene expression during CM maturation.

## Introduction

Aberrant cellular specification and maturation during cardiogenesis may lead to congenital heart disease (CHD)^1^, the most common birth defect worldwide^2^. It may also lead to heart failure, a leading cause of morbidity and mortality in adult life^3^. Since precise control of gene expression is critical for heart development and maturation^4,5^, knowledge of the regulatory mechanisms controlling cardiac gene networks is key to understanding the pathogenesis of cardiovascular disease.

Gene expression is tightly controlled by transcription factors (TFs) that act in accordance with epigenetic regulatory mechanisms^6,7^. TFs bind to specific sequences of DNA adjacent to a gene promoter and control its activity^6^. However, most TFs cannot access closed chromatin, and their binding to DNA relies on the presence of a permissive chromatin environment constructed by epigenetic modulators^7,8^. Global changes in chromatin accessibility have been observed during development of several different species^9,10^, and stage-specific reorganization of the chromatin landscape appears to be critical for gene expression and cell-fate control^11^.

During heart development, early epigenetic marks are established at the embryonic stage^12,13^, and dynamic changes in these marks are associated with cardiac lineage specification and stage transition^13-15^. An increasingly large body of clinical and experimental research indicates that aberrant epigenetic regulation can contribute to CHD^12,16^. For instance, alterations in histone modifying enzymes were shown to commonly occur as *de novo* mutations in CHD patients^16^. More recently, a critical role for histone H2B ubiquitylation (H2Bub) was demonstrated in cardiac development^17,18^. Knockdown of the *Xenopus* H2Bub E3 ligases, *rnf20* (Ring finger protein 20) and *rnf40* (Ring finger protein 40)^19^, beginning at the two-cell stage, influences cardiac looping and left-right (LR) asymmetry during early embryonic development^17^. Furthermore, the results of an *in vivo* CRISPR/Cas9 genetic screen suggested that RNF20/40 is involved in cardiomyocyte (CM) maturation via H2Bub deposition^18^. Thus, RNF20 appears to be relevant to both CHD and CM maturation^17,20^. The addition of a single ubiquitin on H2B disrupts local and higher-order structures of compacted chromatin^21,22^. Through this action, the RNF20-H2Bub pathway regulates gene activities by affecting the binding of TFs or regulatory proteins at gene promoters during transcription^23,24^. H2Bub is also selectively enriched in the coding regions of highly expressed genes^25^, where its tight coupling with RNA polymerase II (Pol II)^23,26^ and histone chaperone FACT^27^ facilitate transcriptional elongation. Nevertheless, the molecular function of RNF20 during temporal development of mammalian heart remains uncharacterized.

Here, we identified an integrated, time-ordered transcriptional network that regulates CM maturation, and we found that deletion of *Rnf20* in CMs disrupts this postnatal gene network and leads to dilated cardiomyopathy. We showed that RNF20 promotes chromatin relaxation in postnatal CMs, playing a key role in the remodeling of chromatin accessibility at this stage. Taken together, our data show how chromatin state transitions can coordinate the temporal expression of distinct transcriptional programs during CM maturation.

## Results

### A time-ordered transcriptional cascade during CM maturation

In postnatal hearts, numerous processes (e.g., cell cycle, structural, functional and metabolic transitions) are required to drive CM maturation^4,5^. These transitional processes are known to be coordinated by certain gene networks^5,29^, but it remains unclear whether the transcriptional programs governing each process are regulated independently and simultaneously or by an integrated time-ordered gene network^4^. To address this open question, we conducted time-series experiments to obtain transcriptomic data from developing hearts (ages P2 to P28) of control mice (Figure 1A). Principal component analysis (PCA) analysis of postnatal heart transcriptomes showed clear distinctive expression profiles among the hearts at different stages (blue circles in Figure 1B), suggesting drastic transcriptional alterations occur during postnatal CM maturation.

**Figure 1.**
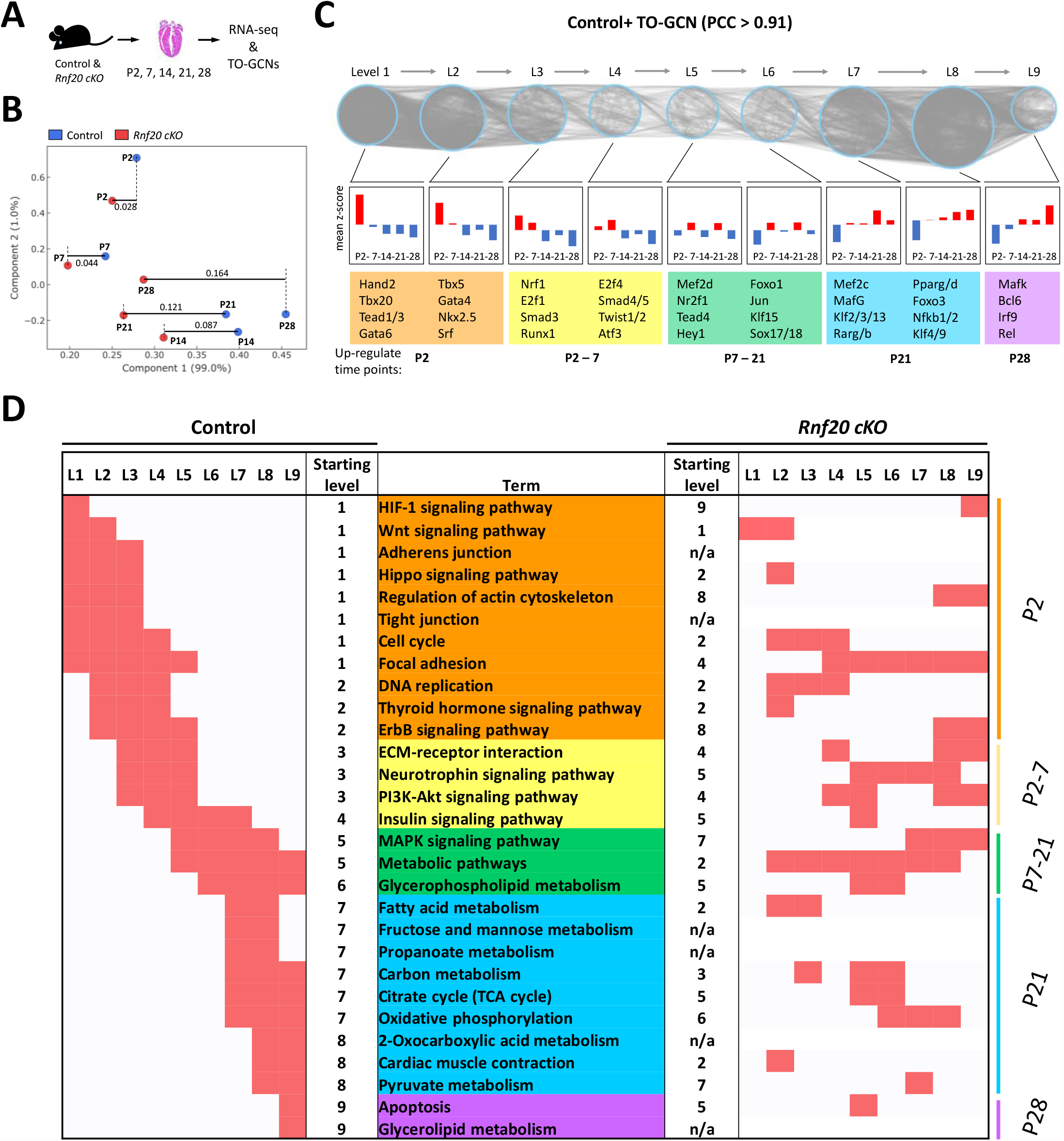
Comparative analysis of time-series transcriptomes from mouse hearts. (A) Schematic of sample collection strategy from control and *cKO* mice for RNA-seq and time-ordered gene co-expression network (TO-GCN) analysis. (B) PCA plot of RNA-seq results. n=2 per group. (C) The TO-GCN structure with TF genes as nodes (blue circles) and the heatmap of average normalized RPKMs (z scores) at each time-point; TF genes were evaluated at nine levels in the control heart. Examples of TFs gene lists from each level in the control mice heart are shown. TFs at L1 and L2 were associated with cardiac development, e.g., Gata4, Nkx2.5, Tbx5, Tbx20 and Tead1, in line with their known contributions to neonatal cell proliferation and maturation^67-69^ (Supplementary file 1). The YAP co-factors (Smad3, Runx1, Smad4/5) were present at L3 and L4 (P7), consistent with their roles in the cardiac homeostasis and cell cycle control at postnatal stages^70^. The postnatal activation of Mef2 family (shown at L5 and L7) is required for normal postnatal growth of the myocardium^71^, and Foxo1 (shown at L6) is necessary for hypertrophic growth^72^. The PPARs shown at L8 are responsible for mitochondrial biogenesis and metabolic switching of CMs^73^. (D) KEGG analysis for co-expressed genes in the control heart are displayed in order (by TO-GCN level); the same list from *cKO* heart is shown for comparison.

The time-ordered gene coexpression network (TO-GCN) algorithm^30^ was applied to elucidate the dynamics of developmental processes (Figure 1C and S1). Each TF that was expressed within the time-series transcriptomes was assigned to one of nine levels (L1 to L9), representing its expression time order over the five time-points (high expression levels are indicated by red squares in Figure 1C and Supplementary file 1). In the network analysis, TFs assigned to the same level are upregulated at the same time period. Moreover, TFs assigned to consecutive levels are upregulated in similar time orders, while TFs at lower levels are upregulated earlier than TFs at higher levels (Figure 1C).

Next, we identified overrepresented functional pathways in heart development at each level of TO-GCN by examining coexpressed genes (Figure 1D and Figure S1-2). The result shows a clear time-ordered organization of transcriptional cascades that regulate CM maturation (Left panel, Figure 1D). For example, hypoxia-inducible factor 1 (HIF1) signaling was enriched at level 1 (L1) and quickly decreased after L2 (Left panel, Figure 1D and S2-3), reflecting the transition of hypoxia to normoxia after birth^31^. Following the decrease in HIF1 signaling, most of the enriched pathways from L1 to L5 (corresponding to the first week) were related to cell-cell connections, including genes supporting adherens junctions, actin cytoskeleton, tight junction focal adhesion, and ECM-receptor interactions (Left panel, Figure 1D and Figure S2). Genes related to cell cycle and DNA replication pathways were activated from L1/L2 (P2) to L4 (P7), reflecting a state of hyperplasia during the first week after birth^32^ (Left panel, Figure 1D). Thyroid (T3) hormone signaling (Left panel, Figure 1D), which is required for CM growth and structural maturation^33^, was enriched from L2 to L4. In addition to the upregulation of T3 signaling, we also observed sequential activation of several other signaling pathways. ErbB signaling was first activated at L2, followed by neurotrophin and PI3K-AKT signaling at L3, insulin signaling at L4 and MAPK signaling at L5 (Left panel, Figure 1D). The combination of ErbB and insulin signaling is thought to promote sarcomere synthesis^34,35^. Notably, cross-talk between T3 and PI3K-AKT is important for myosin heavy chain and titin isoform transitions, and it is further critical for myocardial distensibility and mechanosignaling^36^. Moreover, the induction of neurotrophin signaling is required for neurodevelopment and enhances normal CM Ca^2+^ cycling and contraction^37^. These enriched signaling pathways reflect CMs undergoing structural and functional maturation (indicated by hypertrophy, isoform switch and contraction), accompanied by halting of the cell-cycle. The activated insulin signaling at L4 is important for the following fatty acid metabolism and oxidative phosphorylation at L7, reflecting a metabolism transformation in the CMs^38^. At the same time, genes involved in mitochondrial biogenesis and organization were highly expressed (Figure S2), consistent with idea that maturing CMs undergo a transition of primary energy source from glucose to fatty acids^39^. Lastly, the pathway of cardiac muscle contraction was enhanced at L8 (P21) (Left panel, Figure 1D and Figure S2), in line with enhanced heart function at this stage. In summary, the analysis revealed an integrated, that accurately reflects the known coordination of signaling cascades regulating CM maturation in the first three weeks of life.

### Coordinated postnatal gene networks are regulated by epigenetic factor, RNF20

To investigate how an epigenetic regulator might influence the time-ordered transcriptional network (Control, Figure 1D), *Rnf20* was conditionally ablated in CMs after four chamber formation, a transition point between cardiac morphogenesis and maturation^5,28^. This ablation was accomplished by intercrossing *Rnf20*^*flox/flox*^ with *Mck-Cre* mice (*Rnf20*^*flox/flox*^; *Mck-*Cre; hereafter called *cKO*)^40^ (Figure 1A and S4A). We then collected and analyzed the transcriptomic data of developing hearts from *cKO* mice (Figure 1A) and compared the results with the control data. Remarkably, we observed drastically different transcriptional regulatory cascades in *cKO* hearts (Figure 1D and S3), with disturbances observed as early as at L1 (corresponding to P2). For example, the HIF-1 signaling pathway was enriched at L1 in control hearts and L9 in *cKO* hearts. Intriguingly, the most dramatically dysregulated pathways at L1 in *cKO* hearts are related to actin cytoskeleton, tight junctions and adherens junctions, which contribute to myocyte-myocyte communication^43^ and regulate cell shape (Figure 1D and Figure S2). In addition, delay of ErbB signaling and disruptions in many metabolic networks (i.e., inactivation of fructose/mannose, propanoate and 2-oxocarboxylic acid metabolism, and the early activation of fatty acid metabolism) were also observed in c*KO* (Figure 1D). These disruptions were reflected by the inactivation of genes involved in mitochondrial biogenesis (Figure S2). Thus, RNF20 is critical for the coordination of gene networks during CM maturation as early as the initial postnatal response to the hypoxia-to-normoxia shift.

### Ablation of *Rnf20* after chamber formation leads to dilated cardiomyopathy

*Mck-Cre* ablated *Rnf20* in cardiac and skeletal muscle cells beginning at E13.5^40^ (Figure S4). Notably, the conditional deletion of *Rnf20* before chamber formation by *Mesp1-Cre* (activated at embryonic day 6.5)^44^ (Figure S5A) was embryonically lethal (Table S2), and the mutant embryos displayed defects in heart morphogenesis as early as E9.5 (Figure S5B). In contrast, the depletion of RNF20 by *Mck-Cre* (*cKO*) did not affect the Mendelian ratio of each genotype at postnatal day 1 (P1) (Table S1), but it caused the mice to exhibit significantly lower body weights and sudden death within 8 weeks (P56) of birth (Figure 2A and 2B). At birth, both the control (*flox/flox*) and *cKO* mice had hearts with normal structures, and as expected, the cardiac sizes increased with time (Figure 2C). However, the *cKO* hearts developed a pronounced ventricular chamber dilation after P28 (Figure 2C). Echocardiographic analysis revealed that the left ventricle (LV) systolic function, as measured by fractional shortening (FS) and ejection fraction (EF), was significantly impaired after P28 (Figure 2D). We also detected abnormal electrocardiogram (ECG) patterns in *cKO* mice beginning at P14 (Figure 2E). The depolarization and repolarization of the *cKO* ventricles were disturbed, exhibiting prolonged QRS complex and QT interval (Figure 2E). ECG also revealed that the *cKO* mice developed irregular heartbeat at P28 (Figure S6). Thus, the *cKO* mice appear to suffer from dilated cardiomyopathy^41^ and arrhythmia, which may cause their sudden death. The cardiac defects were not general effects of myopathy since mice with muscle-specific knockout of *Rnf20* (*Rnf20*^*flox/flox*^; *HSA-Cre*^42^) exhibited identical growth curves, survival rates, heart morphologies and cardiac functions compared to control mice at P49 (Figure S7). Taken together, these findings led us to conclude that RNF20-mediated transcriptional network is essential for postnatal cardiac development and function.

**Figure 2.**
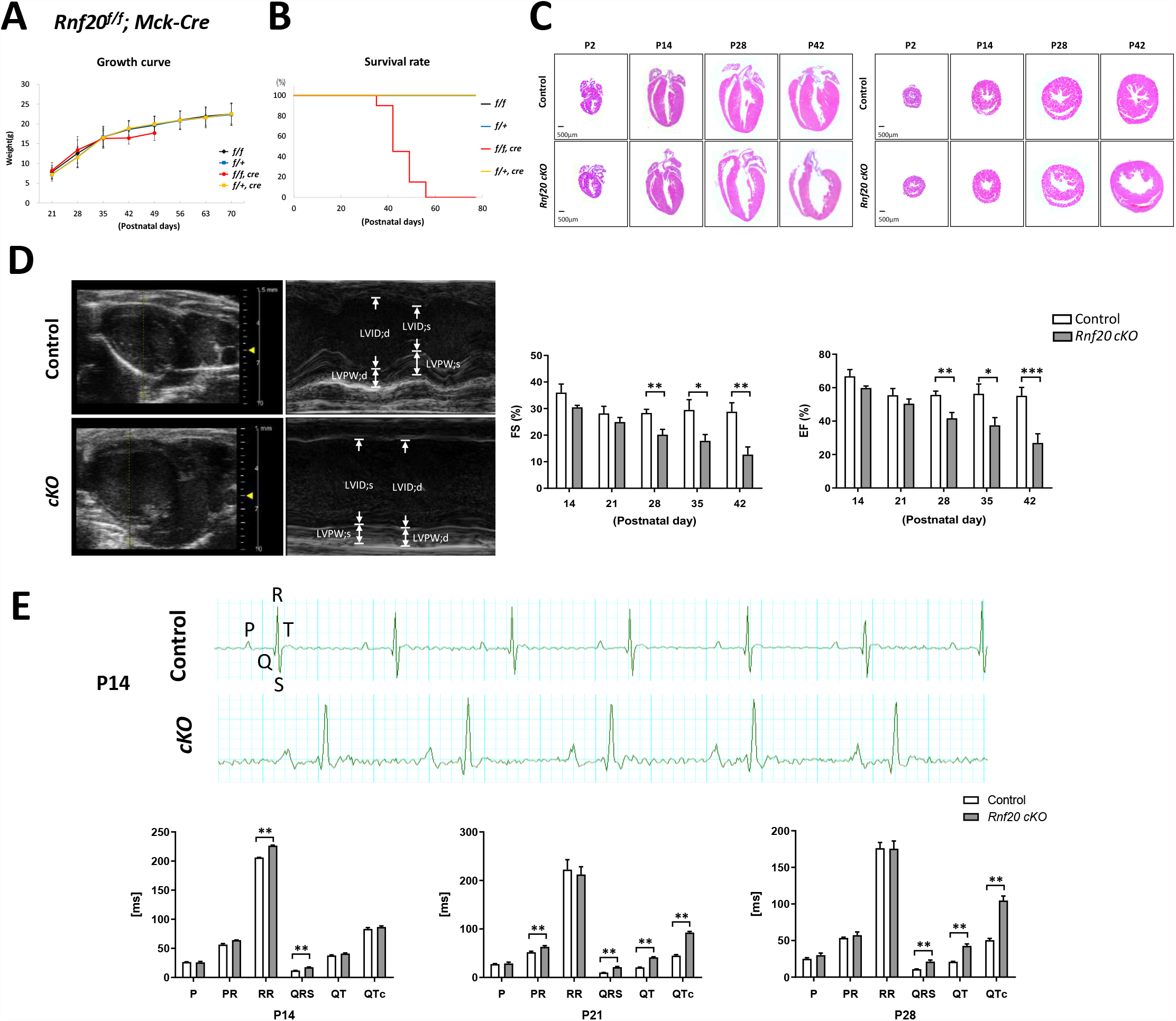
Depletion of RNF20 after chamber formation leads to dilated cardiomyopathy. (A) Growth curve and (B) Kaplan-Meier survival curves of *Rnf20 cKO* (*Rnf20*^*flox/flox*^*;Mck-Cre*) mice and control (*Rnf20*^*flox/flox*^, *Rnf20*^*flox/+*^, *Mck-Cre Rnf20*^*flox/+*^) littermates. (C) Hematoxylin/eosin-stained sagittal (left) and cross (right) sections of control (*Rnf20*^*flox/flox*^) and *Rnf20 cKO* hearts at P2, P14, P28 and P42. (D) (Left panel) Representative 2D and M-mode echocardiographic images of the *RNF20 cKO* and control hearts. (Right panel) Echocardiographic analysis in the control and *cKO* mice at different stages. Left ventricular ejection fraction (EF) and fractional shortening (FS) are shown. Data are presented as mean ± SD; n=6 per group. * p<0.05; ** p< 0.01; *** p< 0.001. (E) Representative telemetric electrocardiogram (ECG) patterns of control and *cKO* mice at P14. Quantification of the ECG changes in *Rnf20 cKO* and control mice from P14 to P42 are shown; data are presented as mean ± SD. n=5 per group. * p<0.05; ** p< 0.01; *** p< 0.001.

### Structural, functional and metabolic transitions in postnatal heart

Since the results up to this point implied that lack of RNF20 may disrupt multiple processes in CM maturation, we sought to analyze the biological outcomes of *Rnf20 cKO* in mouse heart. We first tested whether the loss of RNF20 affects the polarization of intercalated discs (IDs), a hallmark of structural maturation of CMs^5,29^ (Figure 3A). The major components of IDs include N-cadherin (an adherens junction protein), connexin 43 (a gap junction protein) and desmoplakin (a cytoplasmic desmosomal protein). These components were circumferentially distributed at P2 and gradually became polarized (P21), migrating to the ends of the CMs at P35 in the control hearts (Figure 3B and S8A-B). In contrast, the migration was completely abolished in the CMs of *cKO* mice. Next, we characterized the ultrastructural changes in IDs during CM maturation using transmission electron microscopy (TEM). IDs were visible and adjacent to the intercellular space between myocytes in the control CMs at P21 and P35, with clear adherent junctions (white arrow in Figure 3C) and desmosomes (yellow arrow in Figure 3C). By contrast, the structures of IDs in mutant hearts were fragmented and disorganized (P21). By P35, the IDs had become degenerated with widened gaps.

**Figure 3.**
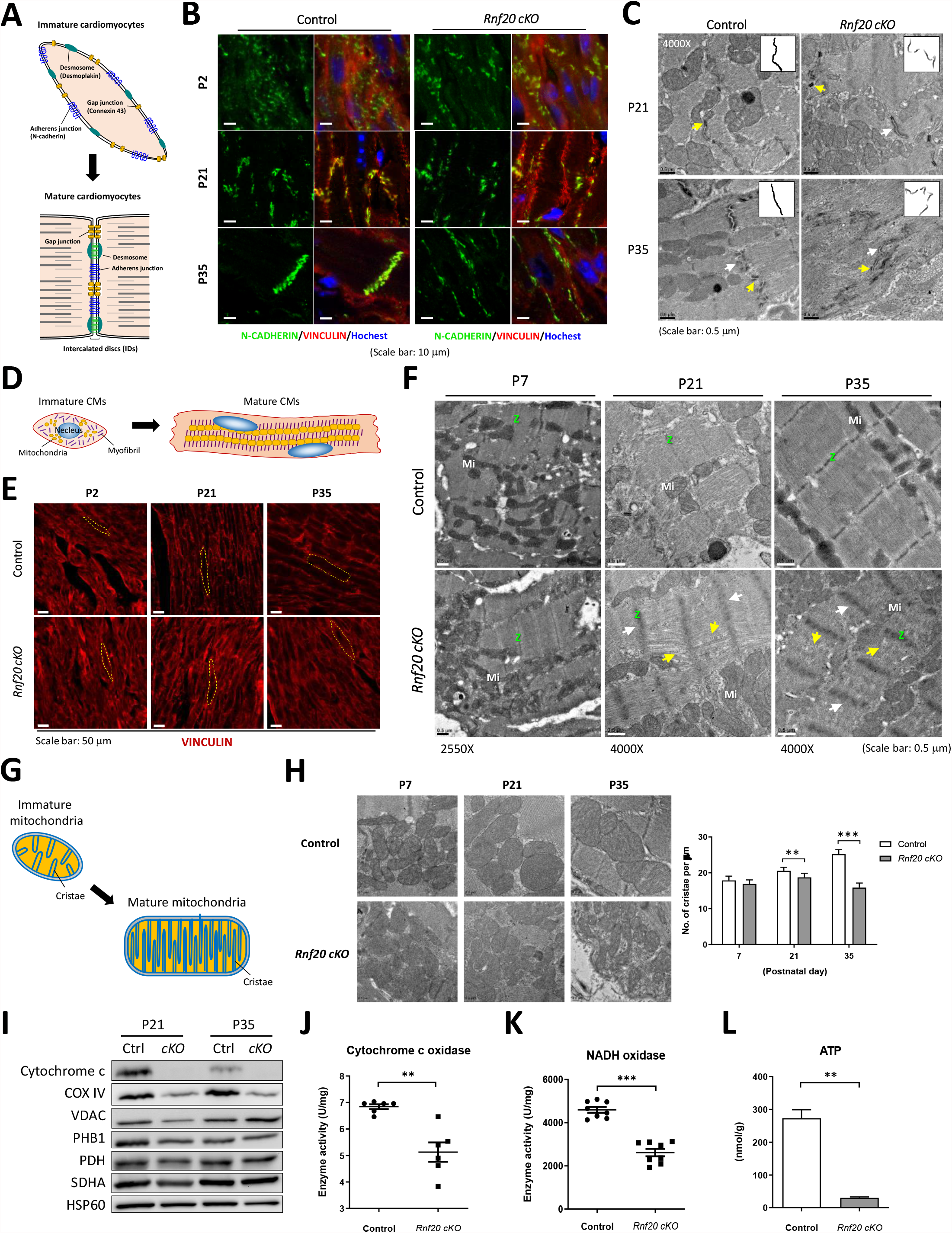
Postnatal maturation defects in *Rnf20 cKO* mice heart. (A) Model of the dynamic formation of intercalated discs (IDs; composed of desmosome, gap junction and adherens junction) during CM maturation. (B) Confocal micrographs of cross sections of the cardiac muscle of control and cKO mice at P2, P21 and P35. Specific antibodies were used to identify the distributions of ID component, N-Cadherin, and the costamere marker, Vinculin. Nuclei were visualized by Hoechst 33342 staining; n=3 per group. Scale bar: 10 μm. (C) Transmission electron microscopy images (TEMs) of ventricular myocardium from *Rnf20 cKO* and control mice at P21 and P35. The insets show representative ID morphologies in control and *cKO* mice. Abnormal structures were observed in mutant hearts, with widened gaps in the IDs. White arrowheads: adherent junctions; yellow arrowheads: desmosomes. Scale bar: 0.5 μm. (D) Model of the morphological changes in immature CMs; increased size and altered organization of the contractile cytoskeleton. (E) Staining for Vinculin (costamere marker) in ventricular sections from *Rnf20 cKO* and control mice at P2, P21 and P35. Scale bar: 50 μm. (F) The ultrastructure of the sarcomeres in CMs of control and *Rnf20 cKO* mice. Abnormal Z-line (white arrowheads) and loose, distorted myofibrils (yellow arrowheads) were observed in *cKO* mice at P21 and P35. Scale bar: 0.5 μm. (G) Schematic depicts the maturation of mitochondria during CM maturation. (H) TEMs of mitochondria in ventricular CMs (Scale bar: 0.5 μm). Quantification of number of cristae per μm suggested a significantly less dense cristae structure in *cKO* mice heart than in control at P35 (*n*LJ=LJ3 for each group). * p<0.05; ** p< 0.01; *** p< 0.001. (I) Immunoblotting for mitochondrial proteins in control and *Rnf20 cKO* hearts (n=3). COX IV: cytochrome c oxidase IV; VDAC: voltage-dependent anion channel; PHB1: prohibitins; PDH: pyruvate dehydrogenase; SDHA: succinate dehydrogenase A; HSP60: heat shock protein 60. (J-L) The mitochondrial enzyme activities for (J) cytochrome c oxidase, (K) NADH oxidase and (L) ATP production were measured in P35 hearts (*n*LJ=LJ3 for each, normalized to mitochondrial protein). * p<0.05; ** p< 0.01; *** p< 0.001.

The structural maturation of CMs (Figure 3D) may also be assessed by the localization of vinculin, a membrane-cytoskeletal protein controlling cytoskeletal mechanics and cell shape^45^ (Figure 3E). In the control mouse myocardium, cytoplasmic vinculin migrated to the cell membrane during maturation (upper panel, Figure 3E); however, the majority of vinculin remained in the cytoplasmic region in RNF20-depleted mutants (lower panel, Figure 3E). Furthermore, we found that the CMs in *cKO* mice failed to adopt a rectangular shape at P35 (Figure 3E and S8C). Since CM cell shape is intimately linked to sarcomere alignment^46^, we also inspected the myofibril organization in postnatal hearts by TEM (Figure 3F). In contrast to the control group, the myofibril alignment in CMs of *cKO* was distorted, exhibiting loose actin filaments (yellow arrow in Figure 3F) and fuzzy Z-lines (white arrowhead in Figure 3F); the M-line was also difficult to distinguish (yellow arrowhead in Figure 3F).

Last, we examined the effects of *Rnf20 cKO* on CM metabolic maturation, as indicated by the maturation of mitochondria (Figure 3G)^4,5^. Control hearts showed a gradual morphological shift from small, tubular and round mitochondria to large, ovoid, well-organized organelles between P7 and P35 (Figure 3H and S8D), which is typical of developing hearts^29^. However, this morphological maturation did not occur in mitochondria of *cKO* hearts (Figure 3H and S8D). We next analyzed the arrangement and density of cristae by TEM. Interestingly, the arrangement of cristae was not altered in *cKO* mutants. However, a breakdown of both inner and outer membrane was observed at postnatal day 35 (*cKO* in Figure 3H). Furthermore, mitochondria cristae density (an indicator of mitochondria function)^5^ was low from P21 onwards in *cKO* hearts (Figure 3H). The mitochondria DNA (mtDNA) copy number was reduced by 20% at P35 in *cKO* mice (Figure S8E). In addition, the protein levels of major electron transport chain components, cytochrome c oxidase and COX VI were markedly decreased in mutant heart (Figure 3I). The observed low enzyme activities of cytochrome c oxidase (Figure 3J) and NADH oxidase (Figure 3K)^47^ were likely responsible for the drastic reduction of ATP in *cKO* myocardium (Figure 3L). Collectively, these data suggested that RNF20 extensively regulates CM maturation by orchestrating gene networks and their functional outcomes.

### The dynamic chromatin opening during CM maturation

Since RNF20 is known to be an epigenetic factor that modulates chromatin compaction via monoubiquitylating histone H2B^21^, we wondered whether Rnf20-mediated epigenetic landscaping is involved in regulating transcriptional networks during postnatal CM development (Figure 1D). To this end, we profiled the chromatin accessibility by ATAC-seq^48^ in postnatal hearts ate early (P7) and late (P21) stages (Figure 4A) of the postnatal transcriptional maturation program. In control hearts, we found the number of wide-spread accessible regions (peaks) was reduced by almost 50% when comparing P21 to P7 (Figure 4B). Most of the diminished peaks were at promoter regions (from 13,320 to 3,714 identified peaks; reduction of 72.1%; Figure 4B and supplementary file 2). Intriguingly, this dynamic change in chromatin accessibility was positively correlated with the level of RNF20 in control hearts (Figure 4C), suggesting that the chromatin in early and late stages displayed distinctive epigenetic landscapes.

**Figure 4.**
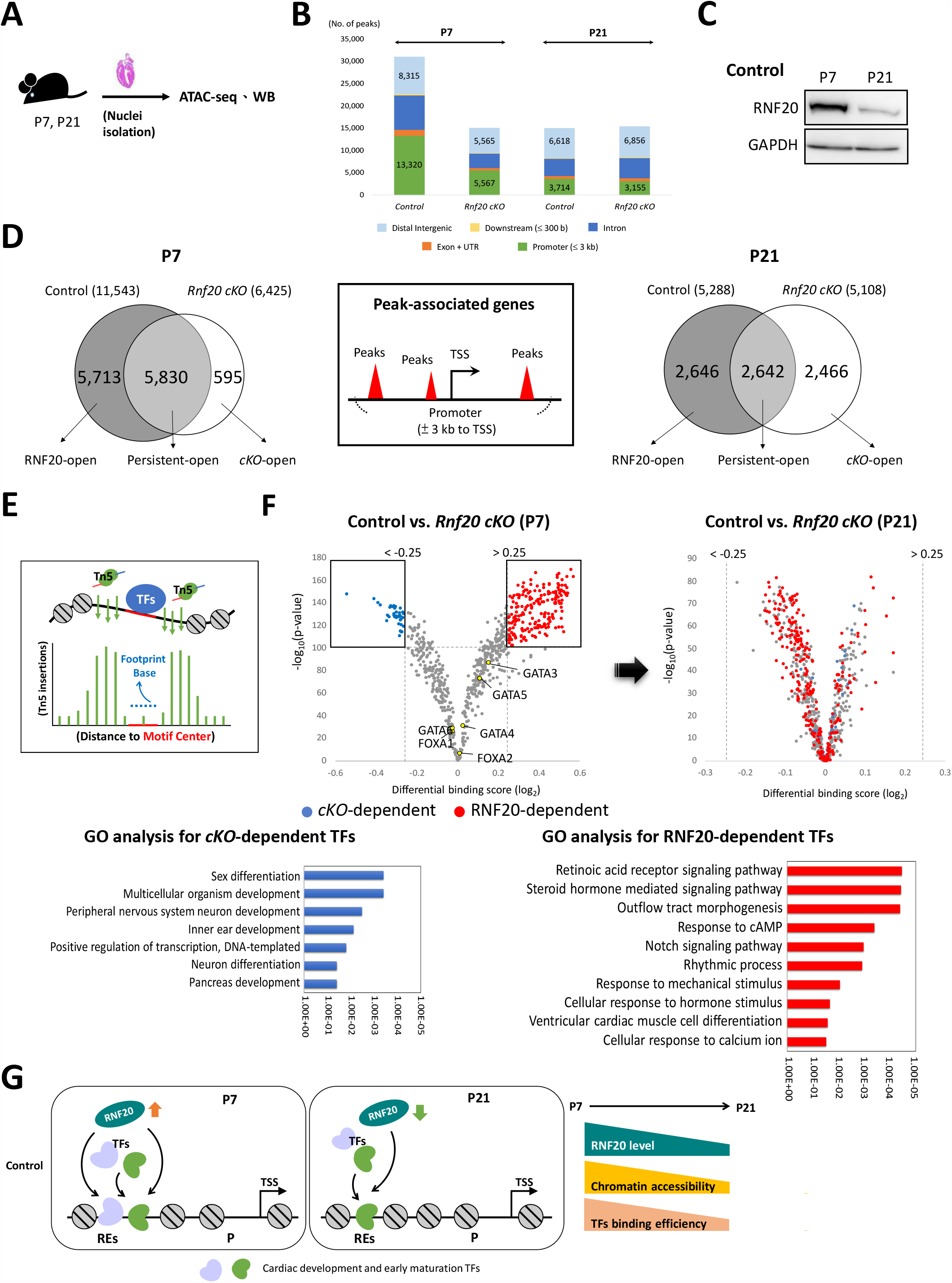
Chromatin accessibility landscape in postnatal heart. (A) Experimental design for sample collection from control and *cKO* mice. (B) Numbers of peak calls from the P7 and P21 control or mutant hearts are shown individually. Color indicates the type of genomic region overlapped by the peak. UTR, untranslated region. (C) Immunoblot analysis of RNF20 in control heart. (D) Venn diagram of peak-associated genes in the control and *Rnf20 cKO* heart at P7 (Left) and P21 (Right), respectively. Middle, model of peak associated genes from ATAC-seq (3kb ± transcription start site). (E) Schematic illustrates the dynamics of TF binding(blue), Tn5 insertion (green), and the concept of footprint analysis using ATAC-seq data. (F) Pairwise comparisons of TF activities in control and *cKO* mice. The volcano plots show differential binding activity versus -log_10_ p-value (both calculated by TOBIAS) for all investigated TF motifs; each dot represents one motif. The control-specific TFs (RNF20-dependent) are labeled in red, and the *cKO*-specific TFs (*cKO*-dependent) are labeled in blue. Gene ontology analyses of RNF20-dependent and *cKO*-dependent TFs are shown. (G) Illustration of RNF20-mediated chromatin remodeling in postnatal heart. P7: RNF20 is upregulated, chromatin accessibility at regulatory elements (REs) of genes are upregulated, TF binding is upregulated. P21: RNF20 is downregulated, chromatin accessibility at REs of genes are downregulated, TF binding is downregulated.

We also found that P7 *cKO* hearts had only about half the number of peaks as P7 controls. The numbers of detected peaks were decreased in all genomic regions but decreases were most predominant in promoter regions (from 13,320 to 5,567 peaks; reduction of 58.2%; Figure 4B). These results suggest that RNF20 facilitates chromatin accessibility, especially during the early stage of postnatal heart development.

### Promoter-opening is required for postnatal gene expression

Since the loss of *Rnf20* primarily impacted accessibility at gene promoters and nucleosome eviction at gene promoter is one of the major events during transcriptional activation^49^, we postulated that RNF20-mediated chromatin opening may affect postnatal transcriptional regulation. To test this hypothesis, we identified genes with ATAC-seq peaks within 3 Kb of the transcription starting sites (TSS) (Figure 4D). A large number of peak-associated genes (11,543 genes) was identified in the control hearts at P7, while only 6,425 genes were associated with accessible promoters in P7 *cKO* hearts (Figure 4D). This indicated that almost half (49.5%; 5,713) of all active genes in the P7 heart may require RNF20 for promoter opening; such genes are hereafter called “RNF20-open” genes (Figure 4D). KEGG pathway enrichment analysis shows that “RNF20-open” genes were enriched in pathways related to CM maturation, such as calcium signaling, cell cycle, fatty acid metabolism, gap junction, and cardiac muscle contraction^4,5^ (Figure S9). Only a small fraction of genes (595 genes) was classified as “*cKO*-open” genes (Figure 4D), indicating their promoters were accessible under RNF20 depletion. Notably, the “*cKO*-open” genes showed no apparent functional links with heart development (Figure S9). These findings revealed that RNF20-mediated chromatin opening at P7 regulates transcriptional programs in CM maturation and thus may play a critical role in coordinating postnatal gene networks.

At P21, the number of peak-associated genes in the control hearts was comparable to that in the *cKO* hearts (5,288 in the control and 5,108 in *cKO*). However, the genes in these two groups overlapped by only about 50% (Figure 4D), and there were roughly equal numbers of “RNF20-open” (2646) and “*cKO*-open” (2466) genes. Of note, the “RNF20-open” genes were involved in metabolism-related pathways, such as pyruvate metabolism, glucagon signaling, regulation of lipolysis in adipocytes (Figure S10). The RNF20-associated pathways mostly correlated with enriched functions at later stages of CM maturation (Figure 1D and S3). In contrast, the “*cKO*-open” genes were enriched in Wnt signaling pathway, ECM-receptor interaction and neurotrophin signaling (Figure S10), which were highly expressed at the neonatal phase (P2 to P7) (Figure 1D). Thus, the ablation of *Rnf20* causes the transcriptional program of P21 heart to retain similarity with the initial postnatal stage and impairs CM maturation.

### Chromatin remodeling facilitates dynamic binding of TFs during postnatal heart development

Previous studies reported that gene network transitions during heart development are tightly controlled by TFs^50^, so we wondered whether RNF20-mediated changes in chromatin accessibility might affect the binding of TFs to modulate postnatal heart development. To answer this question, we used a computational tool, TOBIAS^51^ to perform a TF footprint analysis with the ATAC-seq data (Figure 4E). This tool assigned differential binding scores (DBSs) to 746 TFs, representing the relative binding efficiency between the control and *cKO* groups (Figure 4F; a full list of analyzed TFs can be found in Supplementary file 3). At P7, 199 RNF20-dependent and 39 *cKO*-dependent TFs were respectively identified by their binding activity (-log_10_p-value > 100) in the control (DBS > 0.25) and *cKO* (DBS < -0.25) groups (Figure 4F and supplementary file 3). Gene Ontology (GO) enrichment analysis further showed that the RNF20-dependent TFs were involved in cardiac development and CM maturation (Figure 4F and S11A). Remarkably, many of the RNF20-dependent TFs at P7 turned out to be *cKO*-dependent (DBS < 0) at P21 (Figure 4F), which is in line with the chromatin opening profile in the *cKO* hearts (p21 in Figure 4D). These results provide further explanation for the extensive transcriptional dysregulation caused by deletion of *Rnf20* (Figure 2D). Conversely, the binding patterns of the 39 *cKO*-dependent TFs became sparsely distributed, and their functional categories were unrelated to heart development/function (Figure 4F and S11B). Taken together, these observations suggest that the elevated levels of RNF20 at the early stage (P7) of CM maturation are necessary for proper binding of cardiac TFs that drive transcriptional gene networks, and the continual expression of RNF20 at lower levels is required to sustain gene-network transitions (Figure 4G). Thus, RNF20 is centrally involved in establishing a dynamic epigenetic environment that facilitates the execution of time-ordered gene networks during postnatal heart development (Figure 1D).

### Pioneer factors contribute to chromatin opening

Almost 50% of genes in the control hearts of P7 (50.5%, 5830 genes) and P21 (46.44%, 2642 genes) did not require RNF20 for promoter opening (Figure 4D) and were defined as “persistent-open” genes. The products of “persistent-open” genes included the regulatory factors that are essential for CM maturation, such as HIF-1 signaling, ErbB signaling pathway and tight junctions (Figure 1D, S9 and S10), suggesting there must be other factors that maintain chromatin opening for these maturation aspects. Interestingly, many pioneer factors, including GATA families (GATA4, GATA6) and forkhead box families (FOXA1, FOXA2) (yellow dots in Figure 4F), did not rely on RNF20 for recognition site binding [low Dbs (x-axis) and low p-value (y-axis); Supplementary file 3]. Pioneer factors are able to open closed chromatin sites in order to implement cell fate programs^52^. Consistent with their pioneering roles during embryonic heart development^53,54^, expression levels of GATA4 and GATA6 were high at early postnatal stages (L1-L2 by TO-GCN analysis, Figure 2C). Thus, the promoter accessibilities of “persistent-open” genes are likely attributable to the cooperation of pioneer factors via RNF20-independent mechanisms. Intriguingly, the time-ordered expression of some “persistent-open” genes were disrupted in the absence of RNF20 (Figure 1D, S9 and S10). As such, our data also suggested that RNF20 may help optimize expression of “persistent-open” genes in postnatal heart without affecting chromatin accessibility at their promoters.

## Discussion

In this study, we found that dynamic transitions of chromatin state are required for the time-ordered transcription of postnatal genes and CM maturation. By TO-GCN analysis, we identified a transcriptional regulatory cascade that orchestrates the sequential structural, cell-cycle, functional and metabolic maturation programs in the postnatal heart (Figure 5). Evidence from our study suggests that RNF20 plays a key role in regulating the time-ordered gene networks during postnatal CM maturation by modulating chromatin accessibility. Mechanistically, RNF20-mediated chromatin opening in the early postnatal heart globally resolves epigenetic barriers to allow the maturation program to proceed. The decrease of RNF20 expression at the end-stage of maturation then reestablishes a more compact epigenetic state with less transcriptional flexibility, which is required for the mature cells. RNF20 is likely to modulate chromatin accessibility through its epigenetic target, H2Bub ^27^, as this modification has been shown to disrupt local and higher-order chromatin compaction *in vitro* and *in vivo* ^21,22^. However, our data do not exclude the possibility that RNF20 may regulate other epigenetic factors to affect the postnatal chromatin landscape.

**Figure 5.**
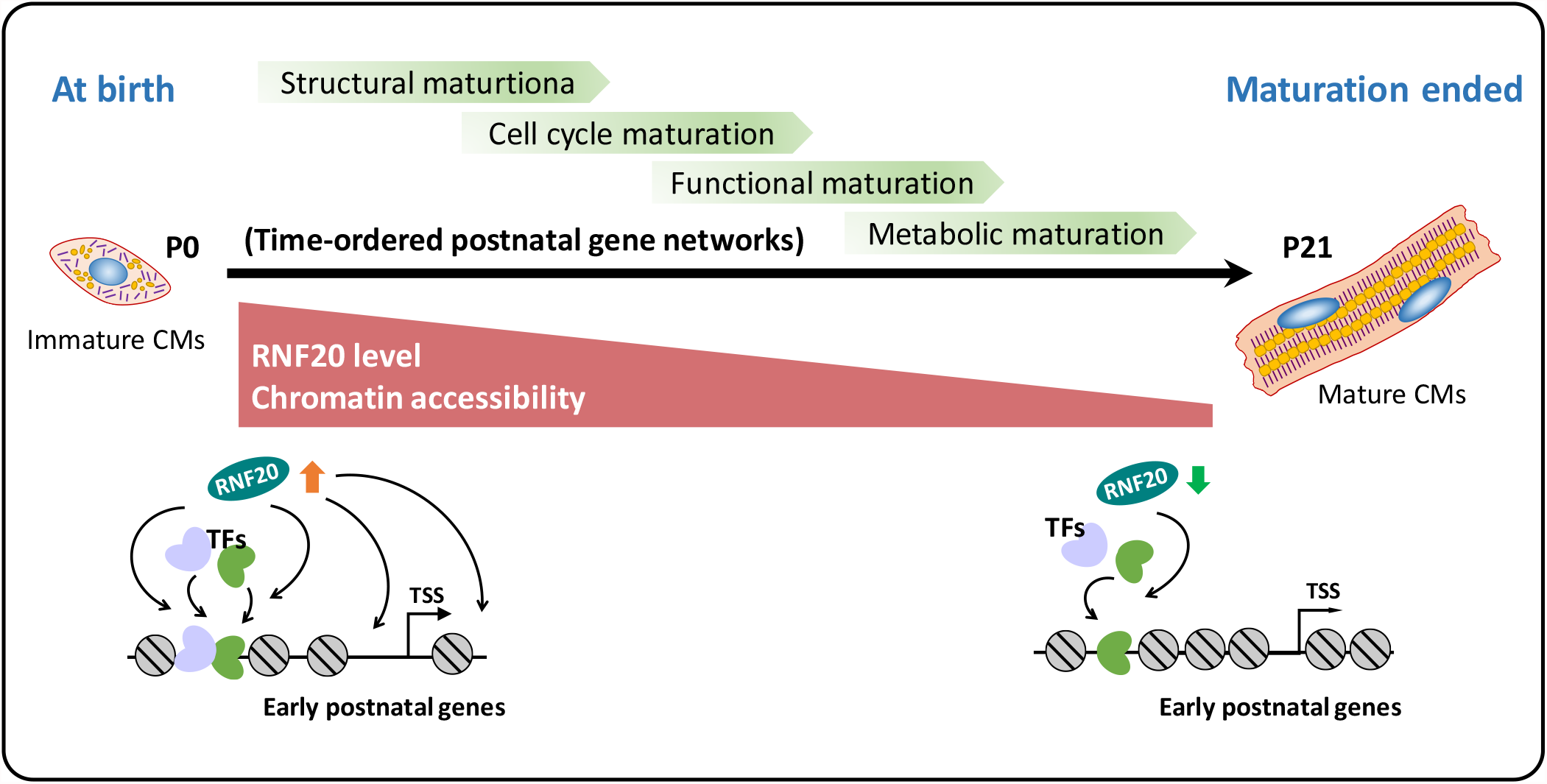
Working model. Dynamic RNF20 expression after birth mediates remodeling of chromatin accessibility to allow the induction of a time-ordered postnatal transcription program. RNF20 is highly expressed at the early postnatal stage, leading to a global chromatin decompaction that facilitates the binding of transcription factors (TFs) to early postnatal genes. RNF20 level decreases at the later stage of heart maturation, leading to chromatin compaction and a restriction of TF binding that establishes and maintains the mature status of the heart.

The dynamics of chromatin accessibility are modulated by the combined effects of chromatin remodelers^56,57^, TFs^58^ and PTMs^59^. However, it remains uncertain how accessibility of chromatin is modulated in response to external stimuli and developmental cues. Overwhelming evidence supports the roles of different PTMs in generating functional chromatin states^60^, and PTMs are known to influence chromatin accessibility via indirect mechanisms, such as altering TF binding and nucleosome affinity of chromatin remodelers. Intriguingly, however, uncertainty remains about how PTMs may directly contribute to accessibility remodeling of chromatin templates. Our finding that RNF20 helps to establish a dynamic chromatin environment for CM maturation has some intriguing implications. First, while dynamic regulation of the RNF20-H2Bub pathway has been linked to embryonic stem cell differentiation^61-63^, the deletion of *Rnf20* after cardiac morphogenesis does not change cell fates or identity during prenatal or neonatal heart development, Instead, it dramatically compromises CM maturation. At the molecular level, we found that RNF20 influences chromatin opening in nearly half of the gene promoters involved in CM maturation, suggesting that there is a division of labor for modulating chromatin accessibility. The cardiac-specific pioneer factors most likely specialize in modulating the chromatin states in those parts of the genome required for cardiac cell fate. In contrast, RNF20 (possibly via H2Bub) may promote chromatin plasticity in the rest of genome, allowing the heart cells to undergo remodeling from a fetal to an adult state without affecting core cell identity factors. Thus, our findings suggest a novel mechanism in which PTMs directly and extensively modulate chromatin accessibility dynamics to orchestrate developmental processes.

The mechanisms of regulating and coordinating constituent processes during CM maturation have received increasing attention because it is not yet possible to achieve complete maturation of pluripotent stem cell (PSC)-induced CMs (iPSC-CMs)^4^. Our TO-GCN analysis revealed that postnatal heart development is initiated by integrated signals, including transient expression of hypoxia-inducible factor 1-alpha (HIF-1α), cell-cell connection (adherens and tight junction) and structural organization (regulation of actin cytoskeleton). Thus, the coordinated activation of these pathways may help initiate CM maturation. In support of this idea, the synergistic application of electrical/mechanical stimulation, extracellular matrix and non-CM co-cultures can facilitate iPSC-CMs maturation^29,61^. Furthermore, the inhibition of HIF1α enhances the contractile, electrophysiologic and metabolic maturation of iPSC-CMs^62,63^. The maturation of CMs involves a complex transcriptional network coordinated by cardiac and non-cardiac cells^29,43,64-66^. Our study provides clear staging information and delineation of transcriptionally active maturation processes based on analyses of whole heart transcriptomes using a comparative transcriptomic method^30^. Thus, our approach may serve to identify the complete set of processes underlying CM maturation. Furthermore, our mouse genetic results imply that RNF20 mutations may cause CHD, as supported by previous observations in CHD patients and in *Xenopus*^16,17^. The results also suggest that functional RNF20 abnormalities beginning at late-gestation periods could be related to early-onset cardiomyopathy in children and young adults, even without observable effects on heart morphogenesis.

In summary, our findings demonstrate that dynamic chromatin accessibility is critical for the time-ordered expression of postnatal genes during CM maturation, and the epigenetic factor RNF20 plays a decisive role in mediating postnatal chromatin accessibility remodeling. We also offer a complete roadmap of gene cascades that direct CM maturation, providing insights that may improve therapies for CHD and facilitate the development of iPSC-CMs to treat heart disease.

## Acknowledgments

The authors thank Dr. Ting-Fen Tsai for constructive comments during manuscript preparation. We are grateful to the staff at the Core Facility of the Institute of Cellular and Organismic Biology, Academia Sinica, for EM service and confocal assistance; the Transgenic Core Facility of the Institute of Molecular Biology, Academia Sinica, for transgenic mice production and Genomics core facility of the Institute of Molecular Biology, Academia Sinica, for NGS service. The authors also thank the Taiwan Mouse Clinic, Academia Sinica, and Taiwan Animal Consortium for the technical support in electrocardiography and high-frequency ultrasound, and Dr. Marcus Calkins for English editing. This work was supported by the intramural funding of the Institute of Cellular and Organismic Biology, Academia Sinica, and by grants from the Ministry of Science and Technology, Taiwan (MOST 109-2320-B-001-017-MY3).

## Author contributions

C.-F.K. and Y.-T.Y. conceived the study. C.-F.K., Y.-T.Y. and C.-C.C. designed the experiments. C.-Y.L., H.-Y.T., Y.-L.S., H.-H.Y. and Y.-R.L. performed the experiments. Y.-M.C. designed and performed the bioinformatic analysis. C.-Y.L. and C.-L.H. analyzed the data. Y.-L.S. and H.-H.Y. provided study materials and animals. C.-Y.L. and C.-F.K. wrote the manuscript.

## Supplementary information

Experimental designs, materials, additional information and references are available in the supplemented files.

## References

1 Epstein, J.A. Franklin H. Epstein Lecture. Cardiac development and implications for heart disease. N Engl J Med 363, 1638–1647, doi:10.1056/NEJMra1003941 (2010).

2 Hoffman, J. I. & Kaplan, S. The incidence of congenital heart disease. J Am Coll Cardiol 39, 1890–1900, doi:10.1016/s0735-1097(02)01886-7 (2002).

3 McMurray, J. J. Clinical practice. Systolic heart failure. N Engl J Med 362, 228–238, doi:10.1056/NEJMcp0909392 (2010).

4 Kannan, S. & Kwon, C. Regulation of cardiomyocyte maturation during critical perinatal window. J Physiol 598, 2941–2956, doi:10.1113/JP276754 (2020).

5 Guo, Y. & Pu, W. T. Cardiomyocyte Maturation: New Phase in Development. Circ Res 126, 1086–1106, doi:10.1161/CIRCRESAHA.119.315862 (2020).

6 Latchman, D. S. Transcription factors: an overview. Int J Biochem Cell Biol 29, 1305–1312, doi:10.1016/s1357-2725(97)00085-x (1997).

7 Liu, L., Jin, G. & Zhou, X. Modeling the relationship of epigenetic modifications to transcription factor binding. Nucleic Acids Res 43, 3873–3885, doi:10.1093/nar/gkv255 (2015).

8 Bannister, A. J. & Kouzarides, T. Regulation of chromatin by histone modifications. Cell Res 21, 381–395, doi:10.1038/cr.2011.22 (2011).

9 Liu, L. et al. An integrated chromatin accessibility and transcriptome landscape of human pre-implantation embryos. Nat Commun 10, 364, doi:10.1038/s41467-018-08244-0 (2019).

10 Thomas, S. et al. Dynamic reprogramming of chromatin accessibility during Drosophila embryo development. Genome Biol 12, R43, doi:10.1186/gb-2011-12-5-r43 (2011).

11 Li, D., Shu, X., Zhu, P. & Pei, D. Chromatin accessibility dynamics during cell fate reprogramming. EMBO Rep 22, e51644, doi:10.15252/embr.202051644 (2021).

12 Jarrell, D. K., Lennon, M. L. & Jacot, J. G. Epigenetics and Mechanobiology in Heart Development and Congenital Heart Disease. Diseases 7, doi:10.3390/diseases7030052 (2019).

13 Paige, S. L. et al. A temporal chromatin signature in human embryonic stem cells identifies regulators of cardiac development. Cell 151, 221–232, doi:10.1016/j.cell.2012.08.027 (2012).

14 Wamstad, J. A. et al. Dynamic and coordinated epigenetic regulation of developmental transitions in the cardiac lineage. Cell 151, 206–220, doi:10.1016/j.cell.2012.07.035 (2012).

15 Gilsbach, R. et al. Dynamic DNA methylation orchestrates cardiomyocyte development, maturation and disease. Nat Commun 5, 5288, doi:10.1038/ncomms6288 (2014).

16 Zaidi, S. et al. De novo mutations in histone-modifying genes in congenital heart disease. Nature 498, 220-+, doi:10.1038/nature12141 (2013).

17 Robson, A. et al. Histone H2B monoubiquitination regulates heart development via epigenetic control of cilia motility. Proc Natl Acad Sci U S A 116, 14049–14054, doi:10.1073/pnas.1808341116 (2019).

18 VanDusen, N. J. et al. Massively parallel in vivo CRISPR screening identifies RNF20/40 as epigenetic regulators of cardiomyocyte maturation. Nat Commun 12, 4442, doi:10.1038/s41467-021-24743-z (2021).

19 Fuchs, G. & Oren, M. Writing and reading H2B monoubiquitylation. Biochim Biophys Acta 1839, 694–701, doi:10.1016/j.bbagrm.2014.01.002 (2014).

20 VanDusen, N. J. et al. Massively parallel in vivo CRISPR screening identifies RNF20/40 as epigenetic regulators of cardiomyocyte maturation. Nat Commun 12, 4442, doi:10.1038/s41467-021-24743-z (2021).

21 Fierz, B. et al. Histone H2B ubiquitylation disrupts local and higher-order chromatin compaction. Nat Chem Biol 7, 113–119, doi:10.1038/nchembio.501 (2011).

22 Moyal, L. et al. Requirement of ATM-dependent monoubiquitylation of histone H2B for timely repair of DNA double-strand breaks. Mol Cell 41, 529–542, doi:10.1016/j.molcel.2011.02.015 (2011).

23 Zhang, F. & Yu, X. WAC, a functional partner of RNF20/40, regulates histone H2B ubiquitination and gene transcription. Mol Cell 41, 384–397, doi:10.1016/j.molcel.2011.01.024 (2011).

24 Kim, J., Hake, S. B. & Roeder, R. G. The human homolog of yeast BRE1 functions as a transcriptional coactivator through direct activator interactions. Mol Cell 20, 759–770, doi:10.1016/j.molcel.2005.11.012 (2005).

25 Minsky, N. et al. Monoubiquitinated H2B is associated with the transcribed region of highly expressed genes in human cells. Nat Cell Biol 10, 483–488, doi:10.1038/ncb1712 (2008).

26 Xiao, T. et al. Histone H2B ubiquitylation is associated with elongating RNA polymerase II. Mol Cell Biol 25, 637–651, doi:10.1128/MCB.25.2.637-651.2005 (2005).

27 Pavri, R. et al. Histone H2B monoubiquitination functions cooperatively with FACT to regulate elongation by RNA polymerase II. Cell 125, 703–717, doi:10.1016/j.cell.2006.04.029 (2006).

28 Krishnan, A. et al. A detailed comparison of mouse and human cardiac development. Pediatr Res 76, 500–507, doi:10.1038/pr.2014.128 (2014).

29 Karbassi, E. et al. Cardiomyocyte maturation: advances in knowledge and implications for regenerative medicine. Nat Rev Cardiol 17, 341–359, doi:10.1038/s41569-019-0331-x (2020).

30 Chang, Y. M. et al. Comparative transcriptomics method to infer gene coexpression networks and its applications to maize and rice leaf transcriptomes. Proceedings of the National Academy of Sciences of the United States of America 116, 3091–3099, doi:10.1073/pnas.1817621116 (2019).

31 Semenza, G. L. Hydroxylation of HIF-1: oxygen sensing at the molecular level. Physiology (Bethesda) 19, 176–182, doi:10.1152/physiol.00001.2004 (2004).

32 Porrello, E. R. et al. Transient regenerative potential of the neonatal mouse heart. Science 331, 1078–1080, doi:10.1126/science.1200708 (2011).

33 Grais, I. M. & Sowers, J. R. Thyroid and the heart. Am J Med 127, 691–698, doi:10.1016/j.amjmed.2014.03.009 (2014).

34 Sanchez-Soria, P. & Camenisch, T. D. ErbB signaling in cardiac development and disease. Semin Cell Dev Biol 21, 929–935, doi:10.1016/j.semcdb.2010.09.011 (2010).

35 Bertrand, L., Horman, S., Beauloye, C. & Vanoverschelde, J. L. Insulin signalling in the heart. Cardiovasc Res 79, 238–248, doi:10.1093/cvr/cvn093 (2008).

36 Kruger, M. et al. Thyroid hormone regulates developmental titin isoform transitions via the phosphatidylinositol-3-kinase/ AKT pathway. Circ Res 102, 439–447, doi:10.1161/CIRCRESAHA.107.162719 (2008).

37 Feng, N. et al. Constitutive BDNF/TrkB signaling is required for normal cardiac contraction and relaxation. Proc Natl Acad Sci U S A 112, 1880–1885, doi:10.1073/pnas.1417949112 (2015).

38 Riehle, C. & Abel, E. D. Insulin Signaling and Heart Failure. Circ Res 118, 1151–1169, doi:10.1161/CIRCRESAHA.116.306206 (2016).

39 Lopaschuk, G. D. & Jaswal, J. S. Energy metabolic phenotype of the cardiomyocyte during development, differentiation, and postnatal maturation. J Cardiovasc Pharmacol 56, 130–140, doi:10.1097/FJC.0b013e3181e74a14 (2010).

40 Johnson, J. E., Wold, B. J. & Hauschka, S. D. Muscle Creatine-Kinase Sequence Elements Regulating Skeletal and Cardiac-Muscle Expression in Transgenic Mice. Molecular and Cellular Biology 9, 3393–3399, doi:Doi 10.1128/Mcb.9.8.3393 (1989).

41 Japp, A. G., Gulati, A., Cook, S. A., Cowie, M. R. & Prasad, S. K. The Diagnosis and Evaluation of Dilated Cardiomyopathy. J Am Coll Cardiol 67, 2996–3010, doi:10.1016/j.jacc.2016.03.590 (2016).

42 Miniou, P. et al. Gene targeting restricted to mouse striated muscle lineage. Nucleic Acids Res 27, e27, doi:10.1093/nar/27.19.e27 (1999).

43 Tirziu, D., Giordano, F. J. & Simons, M. Cell communications in the heart. Circulation 122, 928–937, doi:10.1161/CIRCULATIONAHA.108.847731 (2010).

44 Saga, Y. et al. MesP1: a novel basic helix-loop-helix protein expressed in the nascent mesodermal cells during mouse gastrulation. Development 122, 2769–2778 (1996).

45 Goldmann, W. H. & Ingber, D. E. Intact vinculin protein is required for control of cell shape, cell mechanics, and rac-dependent lamellipodia formation. Biochem Biophys Res Commun 290, 749–755, doi:10.1006/bbrc.2001.6243 (2002).

46 Bray, M. A., Sheehy, S. P. & Parker, K. K. Sarcomere alignment is regulated by myocyte shape. Cell Motil Cytoskeleton 65, 641–651, doi:10.1002/cm.20290 (2008).

47 Anraku, Y. Bacterial electron transport chains. Annu Rev Biochem 57, 101–132, doi:10.1146/annurev.bi.57.070188.000533 (1988).

48 Corces, M. R. et al. An improved ATAC-seq protocol reduces background and enables interrogation of frozen tissues. Nat Methods 14, 959–962, doi:10.1038/nmeth.4396 (2017).

49 John, S. et al. Chromatin accessibility pre-determines glucocorticoid receptor binding patterns. Nat Genet 43, 264–268, doi:10.1038/ng.759 (2011).

50 Uosaki, H. et al. Transcriptional Landscape of Cardiomyocyte Maturation. Cell Rep 13, 1705–1716, doi:10.1016/j.celrep.2015.10.032 (2015).

51 Bentsen, M. et al. ATAC-seq footprinting unravels kinetics of transcription factor binding during zygotic genome activation. Nat Commun 11, 4267, doi:10.1038/s41467-020-18035-1 (2020).

52 Iwafuchi-Doi, M. & Zaret, K. S. Pioneer transcription factors in cell reprogramming. Genes Dev 28, 2679–2692, doi:10.1101/gad.253443.114 (2014).

53 Sharma, A. et al. GATA6 mutations in hiPSCs inform mechanisms for maldevelopment of the heart, pancreas, and diaphragm. Elife 9, doi:10.7554/eLife.53278 (2020).

54 Tremblay, M., Sanchez-Ferras, O. & Bouchard, M. GATA transcription factors in development and disease. Development 145, doi:10.1242/dev.164384 (2018).

55 Janes, J. et al. Chromatin accessibility dynamics across C. elegans development and ageing. Elife 7, doi:10.7554/eLife.37344 (2018).

56 Iurlaro, M. et al. Mammalian SWI/SNF continuously restores local accessibility to chromatin. Nat Genet 53, 279–287, doi:10.1038/s41588-020-00768-w (2021).

57 Schick, S. et al. Acute BAF perturbation causes immediate changes in chromatin accessibility. Nat Genet 53, 269–278, doi:10.1038/s41588-021-00777-3 (2021).

58 Klemm, S. L., Shipony, Z. & Greenleaf, W. J. Chromatin accessibility and the regulatory epigenome. Nat Rev Genet 20, 207–220, doi:10.1038/s41576-018-0089-8 (2019).

59 Allis, C. D. & Jenuwein, T. The molecular hallmarks of epigenetic control. Nat Rev Genet 17, 487–500, doi:10.1038/nrg.2016.59 (2016).

60 Ernst, J. et al. Mapping and analysis of chromatin state dynamics in nine human cell types. Nature 473, 43–49, doi:10.1038/nature09906 (2011).

61 Scuderi, G. J. & Butcher, J. Naturally Engineered Maturation of Cardiomyocytes. Front Cell Dev Biol 5, 50, doi:10.3389/fcell.2017.00050 (2017).

62 Hu, D. et al. Metabolic Maturation of Human Pluripotent Stem Cell-Derived Cardiomyocytes by Inhibition of HIF1alpha and LDHA. Circ Res 123, 1066–1079, doi:10.1161/CIRCRESAHA.118.313249 (2018).

63 Gentillon, C. et al. Targeting HIF-1alpha in combination with PPARalpha activation and postnatal factors promotes the metabolic maturation of human induced pluripotent stem cell-derived cardiomyocytes. J Mol Cell Cardiol 132, 120–135, doi:10.1016/j.yjmcc.2019.05.003 (2019).

64 Pinto, A. R. et al. Revisiting Cardiac Cellular Composition. Circ Res 118, 400–409, doi:10.1161/CIRCRESAHA.115.307778 (2016).

65 Ieda, M. et al. Cardiac fibroblasts regulate myocardial proliferation through beta1 integrin signaling. Dev Cell 16, 233–244, doi:10.1016/j.devcel.2008.12.007 (2009).

66 Brutsaert, D. L. Cardiac endothelial-myocardial signaling: its role in cardiac growth, contractile performance, and rhythmicity. Physiol Rev 83, 59–115, doi:10.1152/physrev.00017.2002 (2003).

67 Liu, R. et al. Tead1 is required for perinatal cardiomyocyte proliferation. PLoS One 14, e0212017, doi:10.1371/journal.pone.0212017 (2019).

68 Pikkarainen, S. et al. GATA-4 is a nuclear mediator of mechanical stretch-activated hypertrophic program. J Biol Chem 278, 23807–23816, doi:10.1074/jbc.M302719200 (2003).

69 Xiang, F. L., Guo, M. & Yutzey, K. E. Overexpression of Tbx20 in Adult Cardiomyocytes Promotes Proliferation and Improves Cardiac Function After Myocardial Infarction. Circulation 133, 1081–1092, doi:10.1161/CIRCULATIONAHA.115.019357 (2016).

70 Wang, J., Liu, S., Heallen, T. & Martin, J. F. The Hippo pathway in the heart: pivotal roles in development, disease, and regeneration. Nat Rev Cardiol 15, 672–684, doi:10.1038/s41569-018-0063-3 (2018).

71 Kolodziejczyk, S. M. et al. MEF2 is upregulated during cardiac hypertrophy and is required for normal post-natal growth of the myocardium. Curr Biol 9, 1203–1206, doi:10.1016/S0960-9822(00)80027-5 (1999).

72 Ferdous, A. et al. FoxO1-Dio2 signaling axis governs cardiomyocyte thyroid hormone metabolism and hypertrophic growth. Nat Commun 11, 2551, doi:10.1038/s41467-020-16345-y (2020).

73 Martin, O. J. et al. A role for peroxisome proliferator-activated receptor gamma coactivator-1 in the control of mitochondrial dynamics during postnatal cardiac growth. Circ Res 114, 626–636, doi:10.1161/CIRCRESAHA.114.302562 (2014).

