## Supplementary Figures for "Chromatin state transition underlies the temporal changes in gene expression during cardiomyocyte maturation"

**Table S1. Mendelian ratios in the P1 stage of *Rnf20*$^{flox/flox}$ and *Rnf20*$^{flox/+}$; *Mck-Cre* mice interbreeding.**

| | Mouse number | Frequency | |
|---|---|---|---|
| Genotype | P1 | Observed | Expected |
| *Rnf20*$^{flox/+}$ | 59 (27M32F) | 24.2% | 25% |
| *Rnf20*$^{flox/flox}$ | 65 (34M31F) | 26.7% | 25% |
| *Rnf20*$^{flox/+}$; *Mck-Cre* | 67 (35M32F) | 27.6% | 25% |
| *Rnf20*$^{flox/flox}$; *Mck-Cre* | 52 (24M28F) | 21.4% | 25% |
| Total (37 litters) | 243 (120M123F) | 100% | 100% |

**Table S2. Mendelian ratios from the *Rnf20*$^{flox/flox}$ and *Rnf20*$^{flox/+}$*;Mesp1-Cre* mice interbreeding.**

| **Age** | Numbers of progeny with the following genotype (%) | | | | | Total number |
|---|---|---|---|---|---|---|
| | **Litters** | ***Rnf20*$^{flox/+}$** | ***Rnf20*$^{flox/flox}$** | ***Rnf20*$^{flox/+}$*;Mesp1-Cre*** | ***Rnf20*$^{flox/flox}$*;Mesp1-Cre*** | |
| **P21** | 4 | 8 (38 %) | 7 (33 %) | 6 (29 %) | 0 (0 %) | 21 |
| **E12.5** | **1** | 1 (11 %) | 1 (11 %) | 4 (44 %) | 3 (33 %) | 9 |
| **E10.5** | **3** | 7 (25 %) | 4 (14 %) | 9 (32 %) | 8 (29 %) | 28 |
| **E9.5** | 6 | 11 (22 %) | 13 (27 %) | 12 (24 %) | 13 (27 %) | 49 |
| **E8.5** | 7 | 17 (29 %) | 16 (27 %) | 14 (24 %) | 12 (20 %) | 59 |

Mice or embryos were isolated at each indicated post-implantation stage and genotyped.

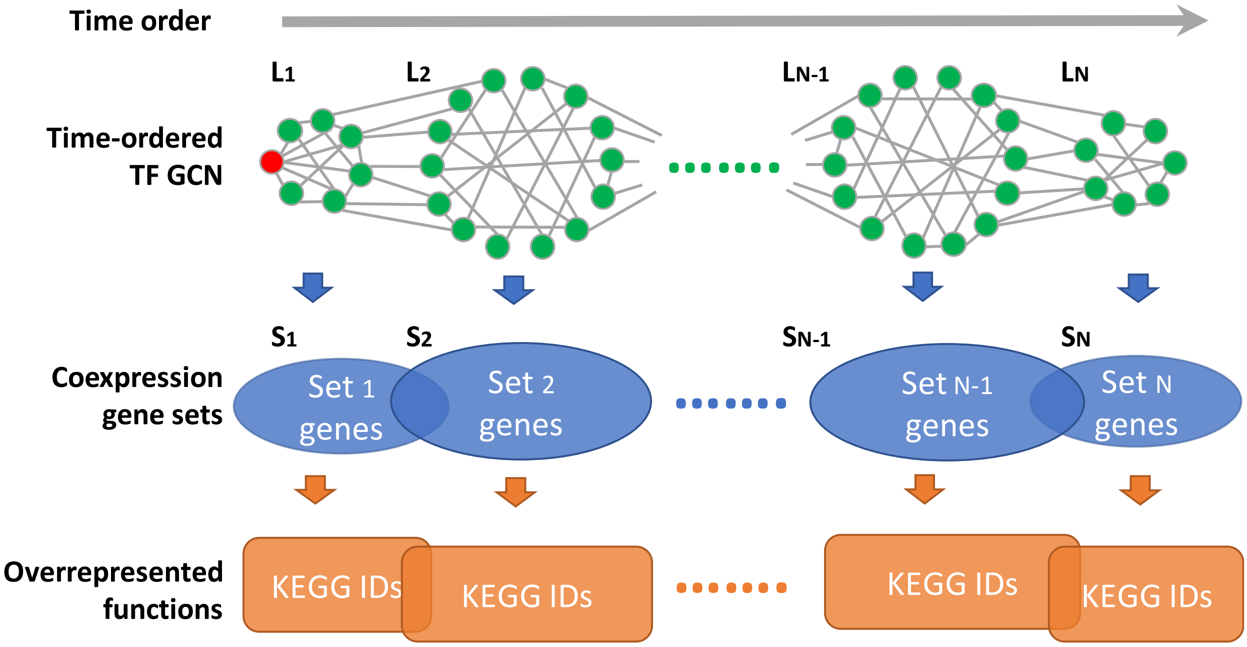

Time order
L1
L2
LN-1
LN
Time-ordered TF GCN
S1
S2
SN-1
SN
Coexpression gene sets
Set 1 genes
Set 2 genes
Set N-1 genes
Set N genes
Overrepresented functions
KEGG IDs
KEGG IDs
KEGG IDs
KEGG IDs


**Figure S1**

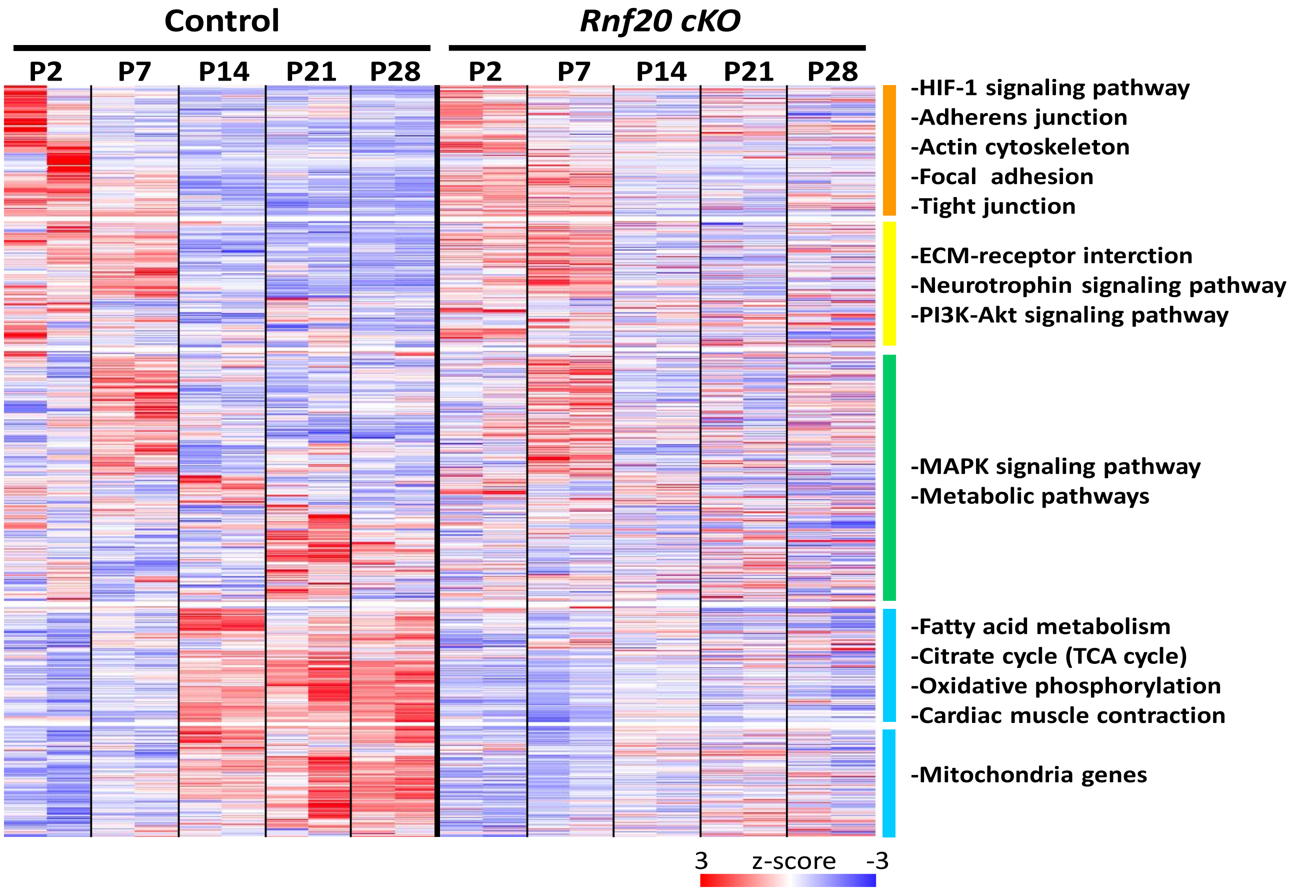

Control
Rnf20 cKO
P2
P7
P14
P21
P28
P2
P7
P14
P21
P28
-HIF-1 signaling pathway
-Adherens junction
-Actin cytoskeleton
-Focal adhesion
-Tight junction
-ECM-receptor interction
-Neurotrophin signaling pathway
-PI3K-Akt signaling pathway
-MAPK signaling pathway
-Metabolic pathways
-Fatty acid metabolism
-Citrate cycle (TCA cycle)
-Oxidative phosphorylation
-Cardiac muscle contraction
-Mitochondria genes
3
z-score
-3


**Figure S2**

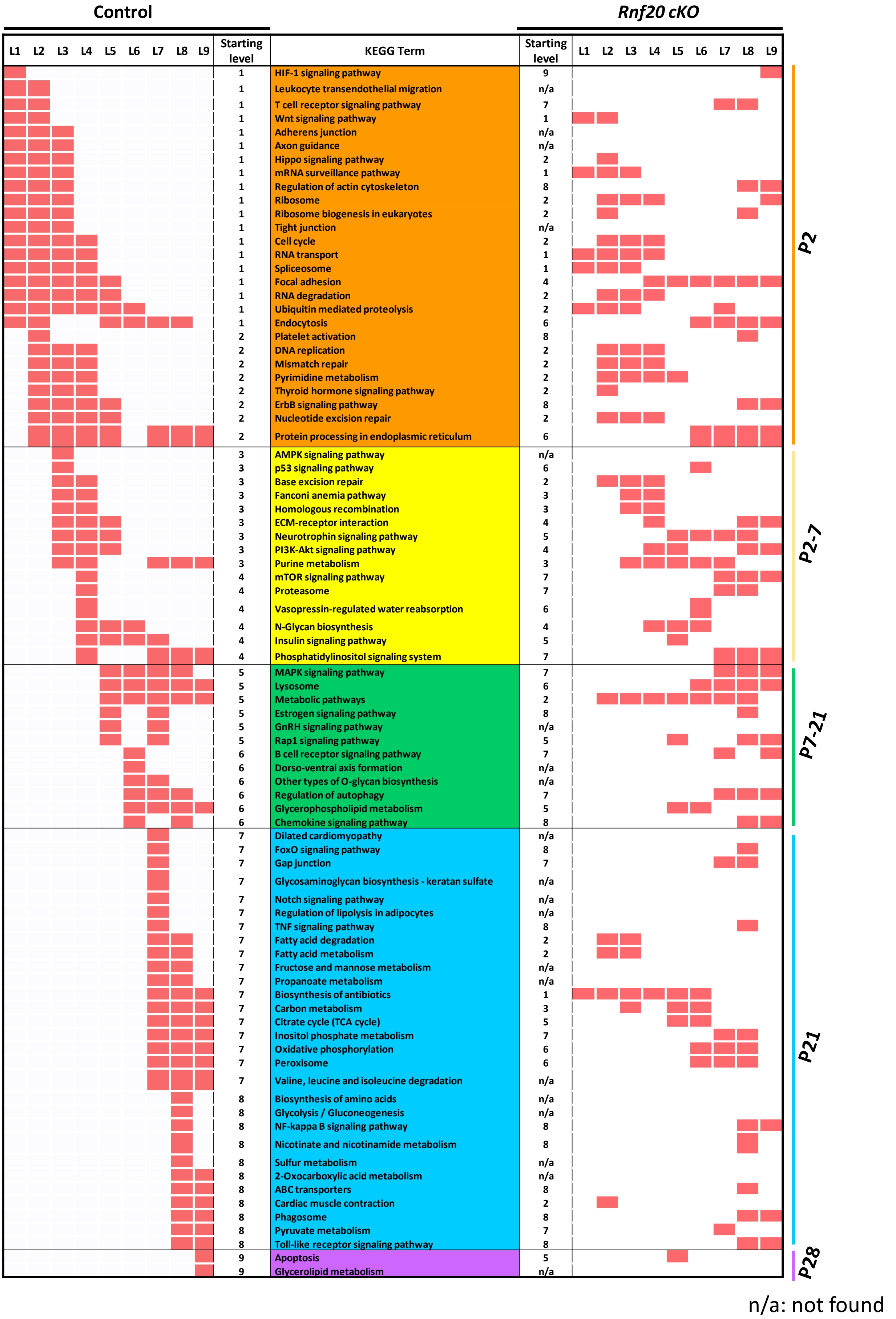

Control
Rnf20 cKO
L1 L2 L3 L4 L5 L6 L7 L8 L9
Starting level
KEGG Term
Starting level
L1 L2 L3 L4 L5 L6 L7 L8 L9
1 HIF-1 signaling pathway 9
1 Leukocyte transendothelial migration n/a
1 T cell receptor signaling pathway 7
1 Wnt signaling pathway 1
1 Adherens junction n/a
1 Axon guidance n/a
1 Hippo signaling pathway 2
1 mRNA surveillance pathway 1
1 Regulation of actin cytoskeleton 8
1 Ribosome 2
1 Ribosome biogenesis in eukaryotes 2
1 Tight junction n/a
1 Cell cycle 2
1 RNA transport 1
1 Spliceosome 1
1 Focal adhesion 4
1 RNA degradation 2
1 Ubiquitin mediated proteolysis 2
1 Endocytosis 6
2 Platelet activation 8
2 DNA replication 2
2 Mismatch repair 2
2 Pyrimidine metabolism 2
2 Thyroid hormone signaling pathway 2
2 ErbB signaling pathway 8
2 Nucleotide excision repair 2
2 Protein processing in endoplasmic reticulum 6
3 AMPK signaling pathway n/a
3 p53 signaling pathway 6
3 Base excision repair 2
3 Fanconi anemia pathway 3
3 Homologous recombination 3
3 ECM-receptor interaction 4
3 Neurotrophin signaling pathway 5
3 PI3K-Akt signaling pathway 4
3 Purine metabolism 3
4 mTOR signaling pathway 7
4 Proteasome 7
4 Vasopressin-regulated water reabsorption 6
4 N-Glycan biosynthesis 4
4 Insulin signaling pathway 5
4 Phosphatidylinositol signaling system 7
5 MAPK signaling pathway 7
5 Lysosome 6
5 Metabolic pathways 2
5 Estrogen signaling pathway 8
5 GnRH signaling pathway n/a
5 Rap1 signaling pathway 5
6 B cell receptor signaling pathway 7
6 Dorso-ventral axis formation n/a
6 Other types of O-glycan biosynthesis n/a
6 Regulation of autophagy 7
6 Glycerophospholipid metabolism 5
6 Chemokine signaling pathway 8
7 Dilated cardiomyopathy n/a
7 FoxO signaling pathway 8
7 Gap junction 7
7 Glycosaminoglycan biosynthesis - keratan sulfate n/a
7 Notch signaling pathway n/a
7 Regulation of lipolysis in adipocytes n/a
7 TNF signaling pathway 8
7 Fatty acid degradation 2
7 Fatty acid metabolism 2
7 Fructose and mannose metabolism n/a
7 Propanoate metabolism n/a
7 Biosynthesis of antibiotics 1
7 Carbon metabolism 3
7 Citrate cycle (TCA cycle) 5
7 Inositol phosphate metabolism 7
7 Oxidative phosphorylation 6
7 Peroxisome 6
7 Valine, leucine and isoleucine degradation n/a
8 Biosynthesis of amino acids n/a
8 Glycolysis / Gluconeogenesis n/a
8 NF-kappa B signaling pathway 8
8 Nicotinate and nicotinamide metabolism 8
8 Sulfur metabolism n/a
8 2-Oxocarboxylic acid metabolism n/a
8 ABC transporters 8
8 Cardiac muscle contraction 2
8 Phagosome 8
8 Pyruvate metabolism 7
8 Toll-like receptor signaling pathway 8
9 Apoptosis 5
9 Glycerolipid metabolism n/a
P2
P2-7
P7-21
P21
P28
n/a: not found


**Figure S3**

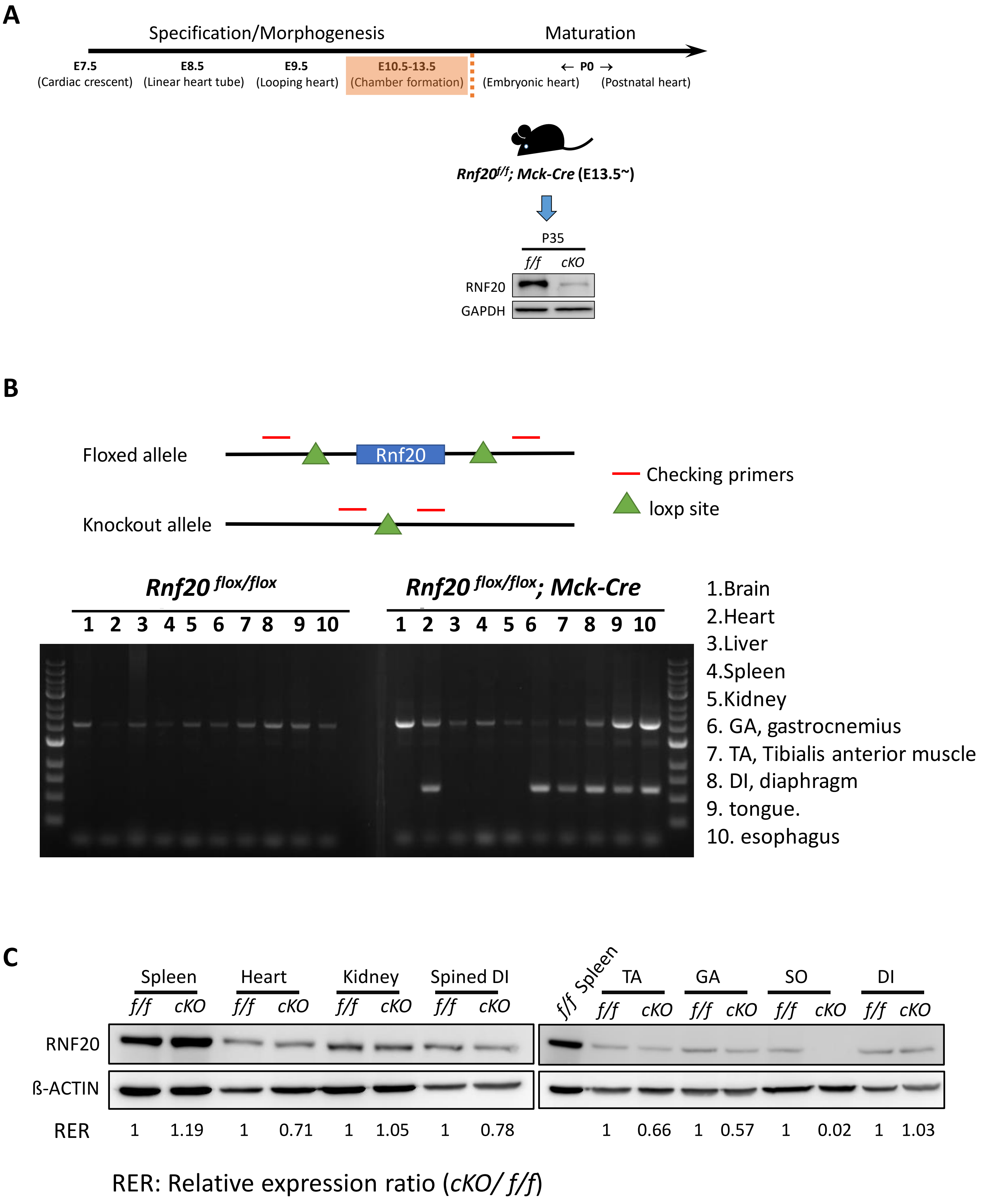
A
Specification/Morphogenesis
Maturation
E7.5
(Cardiac crescent)
E8.5
(Linear heart tube)
E9.5
(Looping heart)
E10.5-13.5
(Chamber formation)
← P0 →
(Embryonic heart)
(Postnatal heart)
Rnf20f/f; Mck-Cre (E13.5~)
P35
f/f cKO
RNF20
GAPDH
B
Floxed allele
Rnf20
Knockout allele
Checking primers
loxp site
Rnf20 flox/flox
Rnf20 flox/flox; Mck-Cre
1 2 3 4 5 6 7 8 9 10
1 2 3 4 5 6 7 8 9 10
1.Brain
2.Heart
3.Liver
4.Spleen
5.Kidney
6. GA, gastrocnemius
7. TA, Tibialis anterior muscle
8. DI, diaphragm
9. tongue.
10. esophagus
C
Spleen
Heart
Kidney
Spined DI
f/f cKO f/f cKO f/f cKO f/f cKO
f/f Spleen
TA
GA
SO
DI
f/f cKO f/f cKO f/f cKO f/f cKO
RNF20
ß-ACTIN
RER 1 1.19 1 0.71 1 1.05 1 0.78 1 0.66 1 0.57 1 0.02 1 1.03
RER: Relative expression ratio (cKO/ f/f)

**Figure S4**

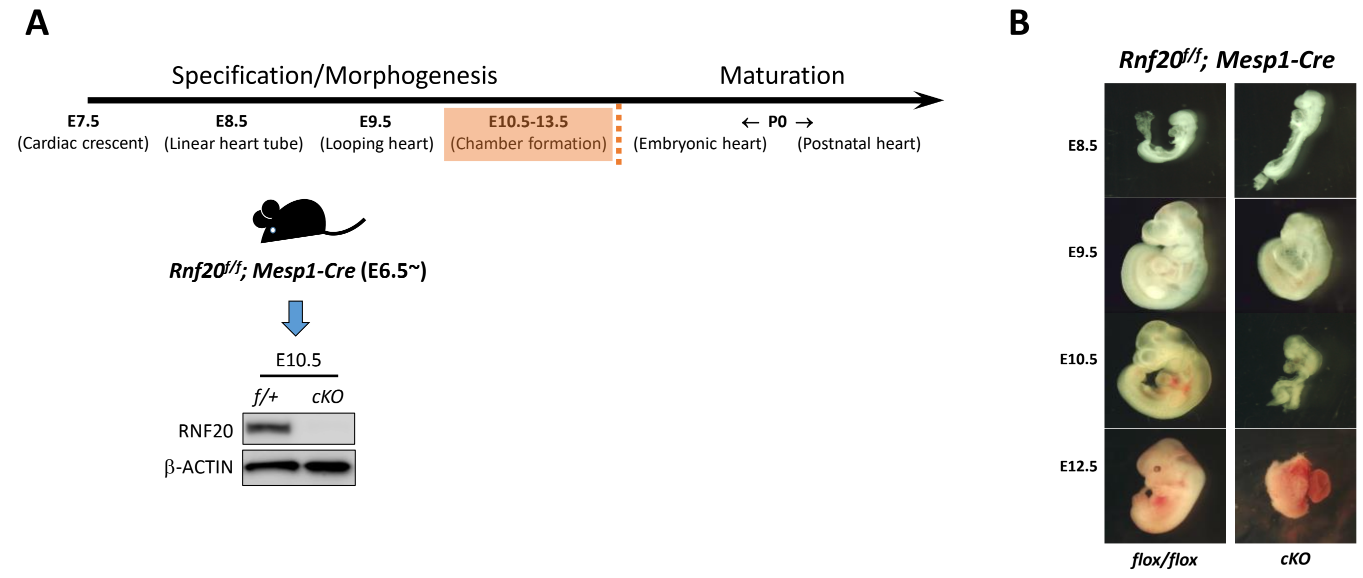
A
Specification/Morphogenesis
Maturation
E7.5
(Cardiac crescent)
E8.5
(Linear heart tube)
E9.5
(Looping heart)
E10.5-13.5
(Chamber formation)
← P0 →
(Embryonic heart)
(Postnatal heart)
Rnf20f/f; Mesp1-Cre (E6.5~)
E10.5
f/+
cKO
RNF20
β-ACTIN
B
Rnf20f/f; Mesp1-Cre
E8.5
E9.5
E10.5
E12.5
flox/flox
cKO

**Figure S5**

# P14

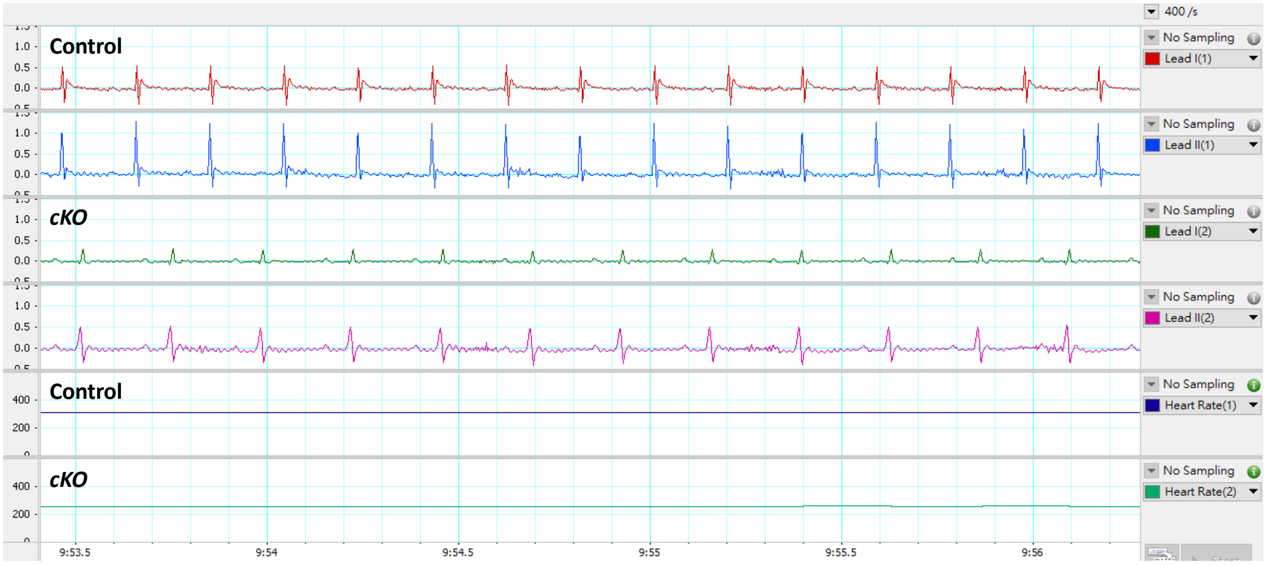


# P28

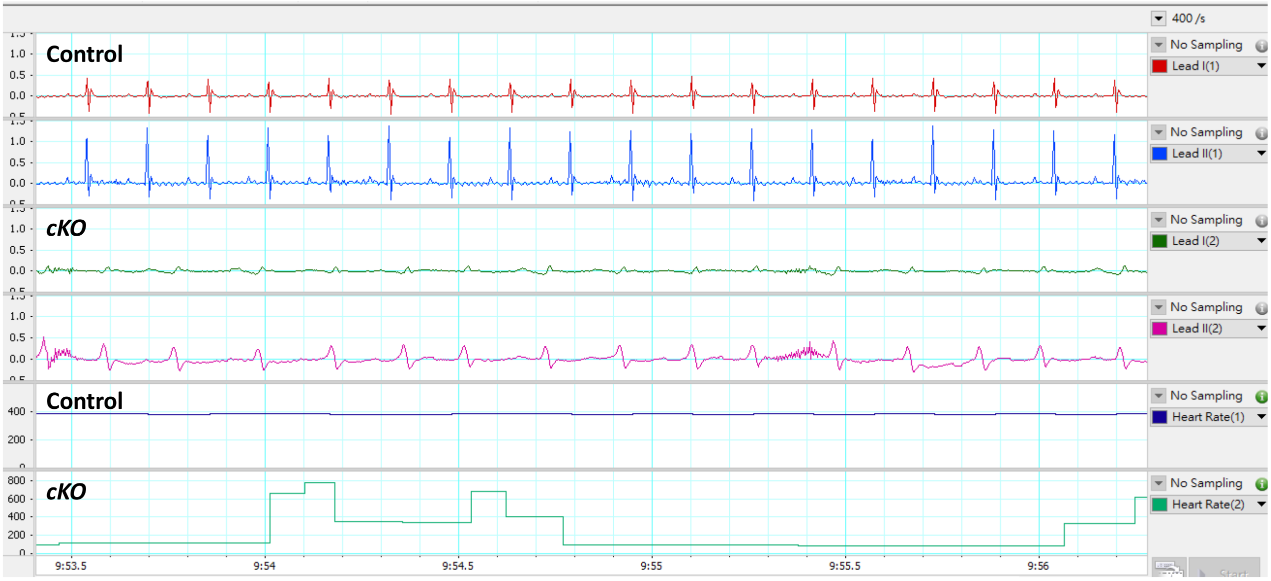


Figure S6

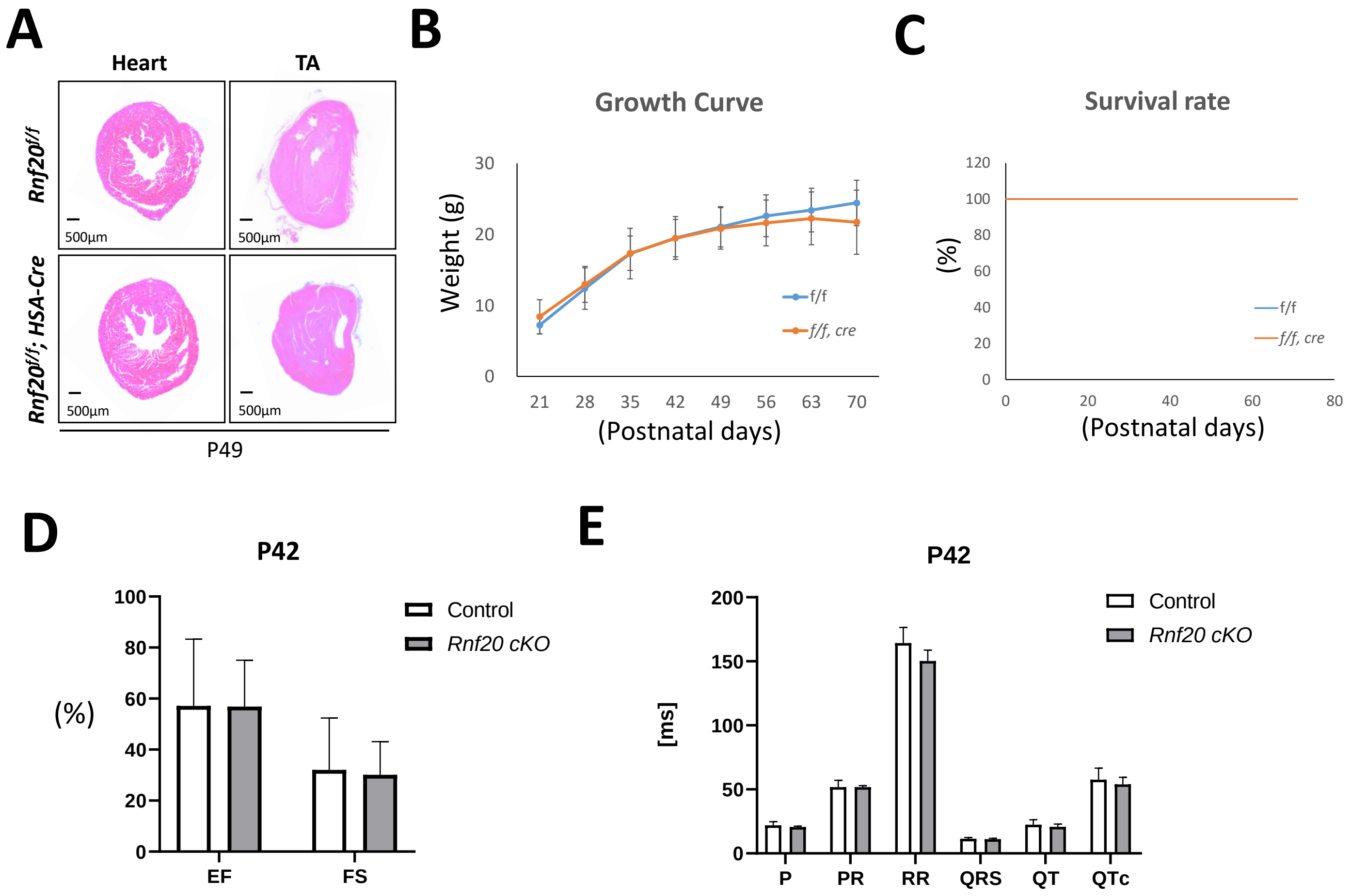
A
Heart
TA
Rnf20f/f
Rnf20f/f; HSA-Cre
500µm
500µm
500µm
500µm
P49
B
Growth Curve
Weight (g)
30
20
10
0
21
28
35
42
49
56
63
70
(Postnatal days)
f/f
f/f, cre
C
Survival rate
(%)
120
100
80
60
40
20
0
0
20
40
60
80
(Postnatal days)
f/f
f/f, cre
D
P42
(%)
100
80
60
40
20
0
EF
FS
Control
Rnf20 cKO
E
P42
[ms]
200
150
100
50
0
P
PR
RR
QRS
QT
QTc
Control
Rnf20 cKO

Figure S7

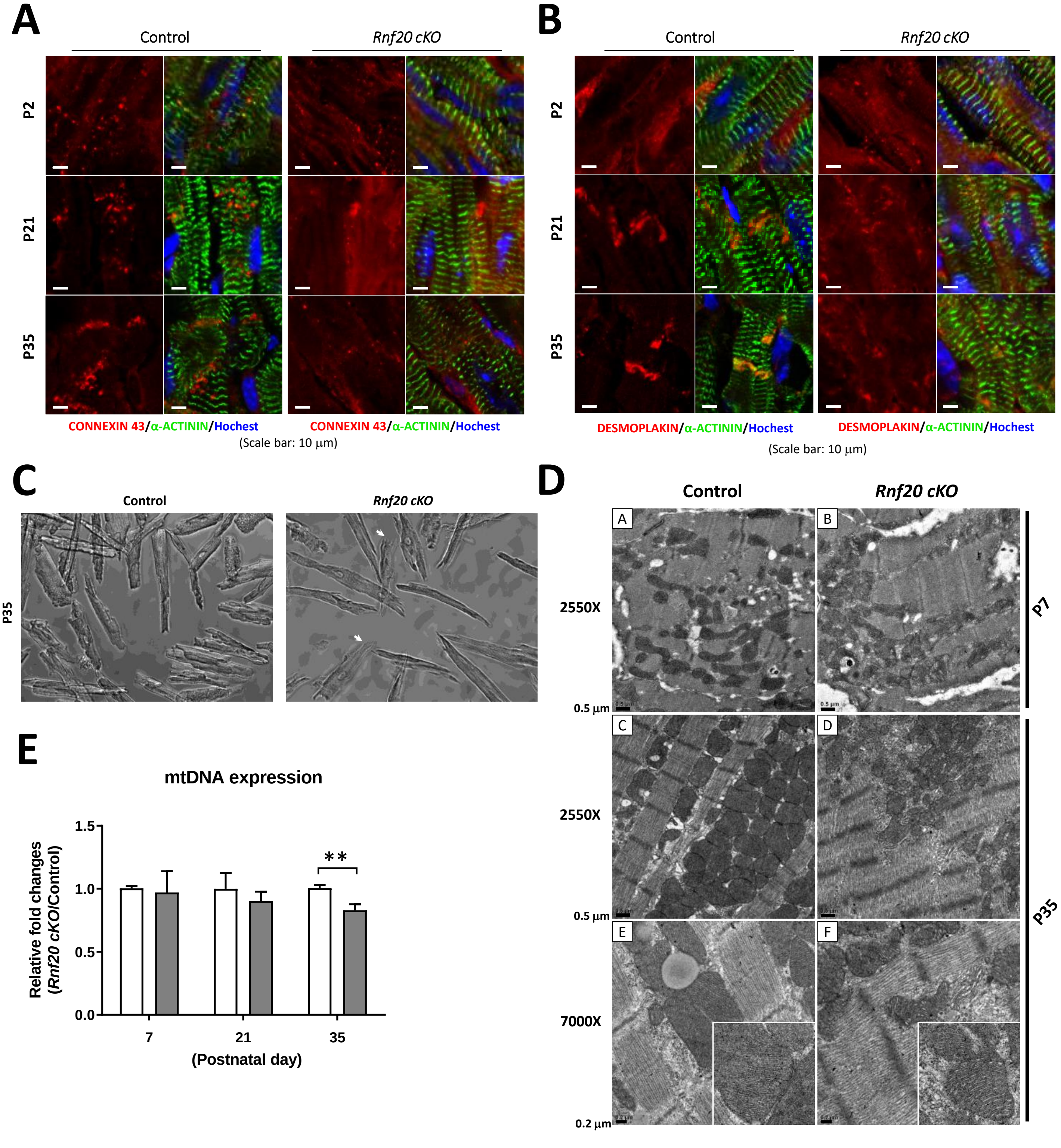
A
Control
Rnf20 cKO
P2
P21
P35
CONNEXIN 43/α-ACTININ/Hochest
CONNEXIN 43/α-ACTININ/Hochest
(Scale bar: 10 μm)
B
Control
Rnf20 cKO
P2
P21
P35
DESMOPLAKIN/α-ACTININ/Hochest
DESMOPLAKIN/α-ACTININ/Hochest
(Scale bar: 10 μm)
C
Control
Rnf20 cKO
P35
D
Control
Rnf20 cKO
A
B
C
D
E
F
2550X
2550X
7000X
0.5 μm
0.5 μm
0.2 μm
P7
P35
E
mtDNA expression
Relative fold changes (Rnf20 cKO/Control)
1.5
1.0
0.5
0.0
**
7
21
35
(Postnatal day)

Figure S8

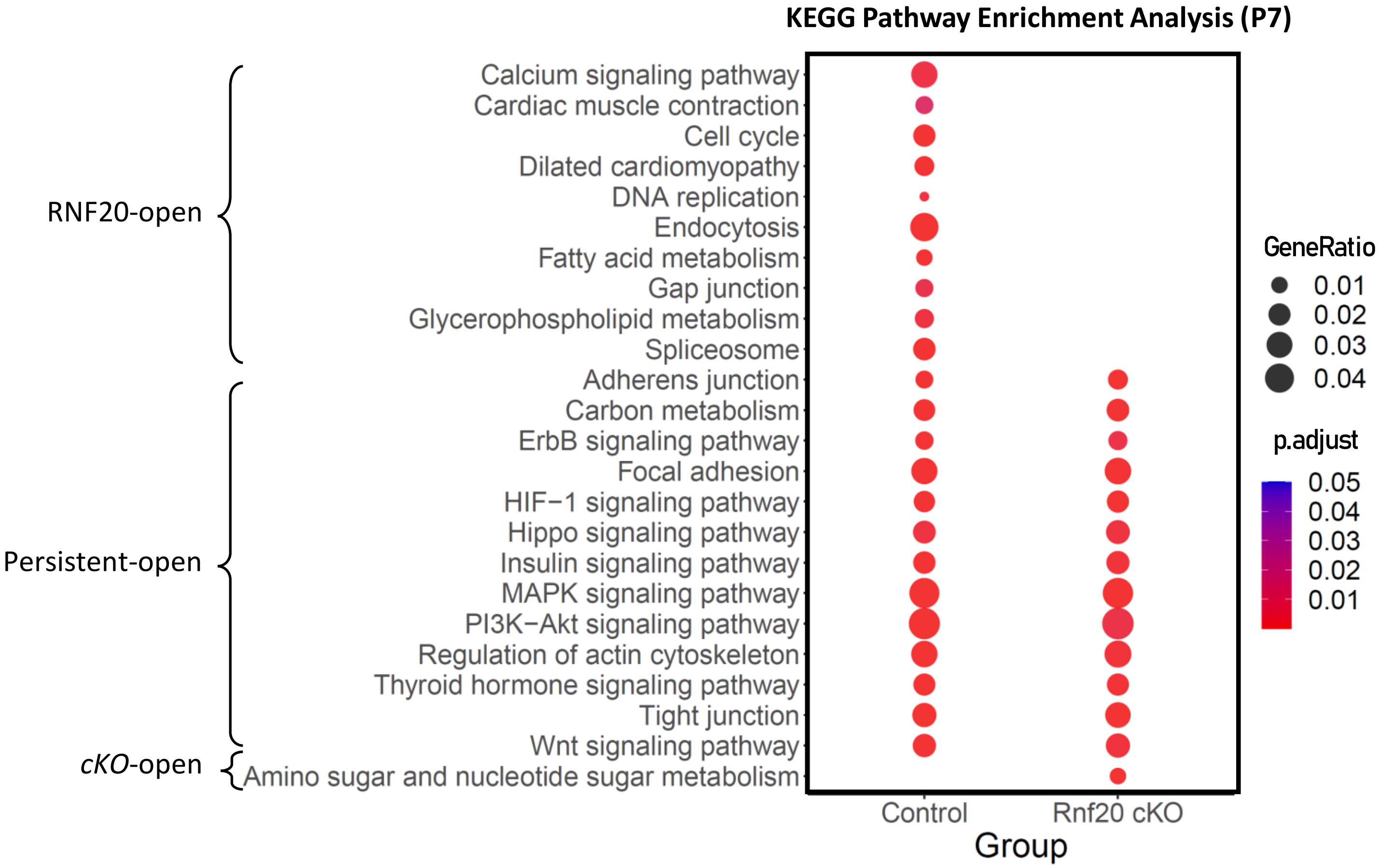
KEGG Pathway Enrichment Analysis (P7)
RNF20-open
Persistent-open
cKO-open
Calcium signaling pathway
Cardiac muscle contraction
Cell cycle
Dilated cardiomyopathy
DNA replication
Endocytosis
Fatty acid metabolism
Gap junction
Glycerophospholipid metabolism
Spliceosome
Adherens junction
Carbon metabolism
ErbB signaling pathway
Focal adhesion
HIF−1 signaling pathway
Hippo signaling pathway
Insulin signaling pathway
MAPK signaling pathway
PI3K−Akt signaling pathway
Regulation of actin cytoskeleton
Thyroid hormone signaling pathway
Tight junction
Wnt signaling pathway
Amino sugar and nucleotide sugar metabolism
Control
Rnf20 cKO
Group
GeneRatio
0.01
0.02
0.03
0.04
p.adjust
0.05
0.04
0.03
0.02
0.01

**Figure S9**

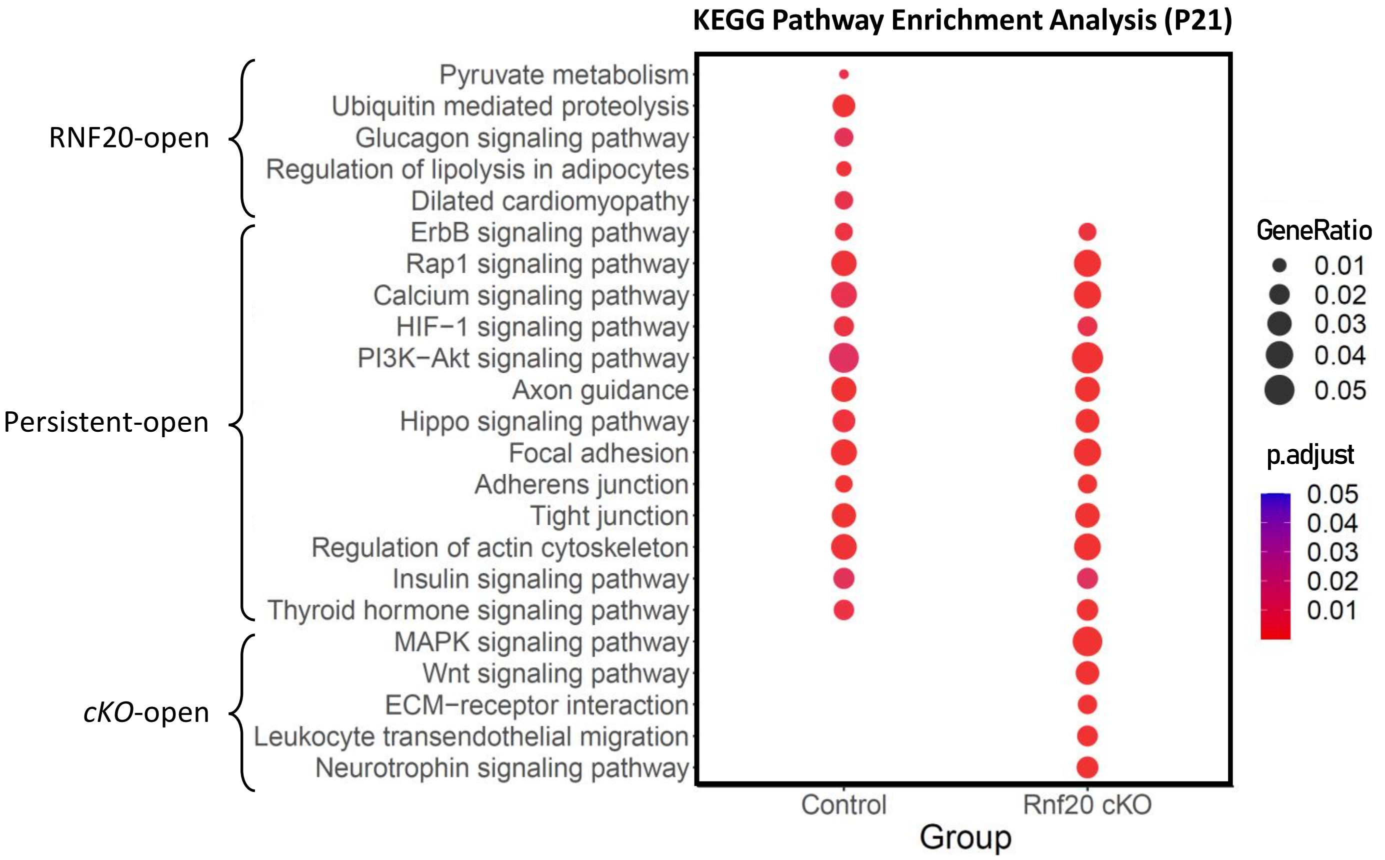
KEGG Pathway Enrichment Analysis (P21)
RNF20-open
Persistent-open
cKO-open
Pyruvate metabolism
Ubiquitin mediated proteolysis
Glucagon signaling pathway
Regulation of lipolysis in adipocytes
Dilated cardiomyopathy
ErbB signaling pathway
Rap1 signaling pathway
Calcium signaling pathway
HIF−1 signaling pathway
PI3K−Akt signaling pathway
Axon guidance
Hippo signaling pathway
Focal adhesion
Adherens junction
Tight junction
Regulation of actin cytoskeleton
Insulin signaling pathway
Thyroid hormone signaling pathway
MAPK signaling pathway
Wnt signaling pathway
ECM−receptor interaction
Leukocyte transendothelial migration
Neurotrophin signaling pathway
Control
Rnf20 cKO
Group
GeneRatio
0.01
0.02
0.03
0.04
0.05
p.adjust
0.05
0.04
0.03
0.02
0.01

**Figure S10**

# A

### GO analysis for RNF20-dependent TFs

| Term | Description | Genes |
|---|---|---|
| GO:0048384 | Retinoic acid receptor signaling pathway | RXRB, RXRA, CREM, RARB, PTF1A, RXRG |
| GO:0043401 | Steroid hormone mediated signaling pathway | NR4A2, RXRB, NR5A1, RXRA, NR2F1, RARB, NR2C2, RXRG |
| GO:0055012 | Ventricular cardiac muscle cell differentiation | RXRB, RXRA, RARB |
| GO:0003151 | Outflow tract morphogenesis | TFAP2A, ATF2, JUN, HEY1, HEY2, HES1, RBPJ, HIF1A |
| GO:0048511 | Rhythmic process | KLF10, HES7, SP1, BHLHE40, BHLHE41, CREM, NPAS2, NFKB2, ARNTL |
| GO:0007219 | Notch signaling pathway | HES7, HEY1, HEY2, HES1, ETV2, RBPJ, ASCL1, ATOH1, HES5 |
| GO:0051591 | Response to cAMP | FOSL1, JUN, JUND, CREM, FOSB, FOS, JUNB |
| GO:0009612 | Response to mechanical stimulus | FOSL1, JUN, JUND, FOSB, JUNB, ETS1 |
| GO:0032870 | Cellular response to hormone stimulus | JUN, JUND, FOSB, FOS, JUNB |
| GO:0071277 | Cellular response to calcium ion | JUN, JUND, FOSB, FOS, JUNB |

GO (BP) enrichment test by DAVID v6.8 with FDR < 0.05

# B

### GO analysis for *cKO*-dependent TFs

| Term | Description | Genes |
|---|---|---|
| GO:0007548 | Sex differentiation | DMRTA2, DMRT1, DMRTC2, DMRT3 |
| GO:0007275 | Multicellular organism development | HOXA10, CEBPA, DMRT1, DMRT3, HOXB13, POU4F1, HOXD9, POU3F3, PHOX2B, POU5F1, PHOX2A |
| GO:0048935 | Peripheral nervous system neuron development | ONECUT2, POU4F1, HOXD9 |
| GO:0048839 | Inner ear development | CEBPA, PHOX2B, POU3F4, POU4F3 |
| GO:0045893 | Positive regulation of transcription, DNA-template | CEBPA, POU1F1, DDIT3, POU2F2, POU3F1, POU3F3, POU5F1 |
| GO:0030182 | Neuron differentiation | POU3F2, POU4F1, POU4F2, PHOX2B |
| GO:0031016 | Pancreas development | ONECUT2, ONECUT1, NKX3-2 |

GO (BP) enrichment test by DAVID v6.8 with FDR < 0.05

Figure S11
