## Supplementary figure legends for "Chromatin state transition underlies the temporal changes in gene expression during cardiomyocyte maturation"

**Figure S1. Comparative transcriptomics method.**

The levels (L_1_ to L_N_) in the TO-GCN analysis represent the upregulation timing of TF genes; the co-expressed gene sets (S_1_ to S_N_) (including non-TF genes) correspond to different levels, and the overrepresented functions. The red-node in L_1_ represents the initial node. Genes in a set may be co-expressed with TFs in multiple levels, so the gene may belong to multiple sets.

**Figure S2. Clustered heatmaps with Z scores for the genes in selected KEGG pathways from the control and *cKO* heart across five postnatal stages.** n=2 per group.

**Figure S3. Full list of KEGG pathways for co-expressed genes in the control heart (by TO-GCN levels) compared with the same lists from *cKO* heart.**

**Figure S4. Characterization of cardiac and muscle-specific RNF20 knockout mouse model by PCR and immunoblotting.**

**(A)** Experimental design for the generation and identification of heart-specific *Rnf20* conditional knockout mice, *Rnf20^flox/flox^; Mck-Cre*. Protein extracts were collected from the isolated cardiomyocytes from *Rnf20^flox/flox^;Mck-Cre* mice (at P35); immunoblot analyses were performed. **(B)** Representative examples of genotyping results are shown for the RNF20 floxed and knockout alleles. Genomic DNA of different tissues was collected from control (*Rnf20^flox/flox^*) or *cKO* (*Rnf20^flox/flox^; Mck-Cre*) mice at P28, and the target regions were amplified using appropriate primers (marked by red line). **(C)** Protein samples were collected at P28 and the RNF20 levels in different tissues were detected by immunoblot analysis using β-actin as loading control (relative expression ratios are shown). (n=2 per group) GA: gastrocnemius; TA: tibialis anterior muscle; SO: soleus; DI: diaphragm.

**Figure S5. Cardiac-specific RNF20 knockout by *Mesp1-Cre* mice.**

**(A)** Experimental design for the generation and identification of heart-specific *Rnf20* conditional knockout mice, *Rnf20^flox/flox^;Mesp1-Cre*. Protein extracts were collected from the heart tube of E10.5 embryos, and immunoblot analyses were performed. **(B)** Bright-field images of *Rnf20^flox/flox^;Mesp1-Cre* embryos and wild-type (*Rnf20^flox/flox^)* littermates at E8.5, 9.5, 10.5 and 12.5.

**Figure S6. Representative telemetric 2-lead ECG.**

Three-second ECG traces indicated arrhythmias in *cKO* (*Rnf20^flox/flox^; Mck-Cre*) mice at P28. (n=5 per group)

**Figure S7. Knockout of *Rnf20* in muscle-specific cre mice (human alpha-skeletal actin (HSA) *-Cre*) does not affect growth or heart function after birth.**

**(A)** Hematoxylin/eosin-stained cross-section of the heart and longitudinal section of the tibialis anterior muscle in *Rnf20^flox/flox^* and *Rnf20^flox/flox^; Hsa-Cre* mice at P49. **(B)** Growth curve and **(C)** Kaplan-Meier survival curves of *Rnf20^flox/flox^* and *Rnf20^flox/flox^;Hsa-Cre* mice. **(D)** Echocardiographic analysis in the control (*Rnf20^flox/flox^*) and *Rnf20 cKO* (*Rnf20^flox/flox^;Hsa-Cre*) mice at P42. Left ventricular ejection fraction (EF) and fractional shortening (FS) are shown. Data are presented as mean ± SD; n=3 per group. **(E)** Quantitative results of the ECG changes in *Rnf20 cKO* and control mice at P42 are shown; data are presented as mean ± SD. n=5 per group.

**Figure S8. Maturation defects in *cKO* mice.**

**(A and B)** Confocal micrographs of cross sections of the cardiac muscle of the control and *cKO* mice at P2, P21 and P35. Specific antibodies were used to identify the distributions of intercalated disc component, Connexin 43 **(A)**, Desmoplakin **(B)** and the sarcomere component, α-Actinin. The nuclei were visualized by Hoechst 33342 staining. n=3 per group. Scale bar: 10 μm. **(C)** Isolation of ventricular cardiomyocytes from the control and *Rnf20 cKO* mice. **(D)** Representative transmission electron micrographs (TEM) sections of the control and *cKO* hearts at P7 and P35 (n=3 per group). Insets show higher magnification views of the mitochondria at P35. Scale bar: 0.5 μm or 0.2 μm. **(E)** Quantitative PCR analysis showed that mitochondrial DNA copy number normalized to nuclear DNA copy number (mtDN1 versus H19 or mtDN2 versus Mx1) was significantly decreased in *RNF20*-depleted heart (*n* = 3 per group).

**Figure S9. Gene set enrichment analysis of the peak-associated genes in the control and *cKO* mice at P7.**

Plot shows the upregulated KEGG pathways (p.adjust <0.05) identified using DAVID as enriched in different groups. The size of the dot is based on gene ratio (gene counts) enriched in the pathway, and the color of the dot shows the pathway enrichment significance.

**Figure S10. Gene set enrichment analysis of the peak-associated genes in the control and *cKO* mice at P21.**

Dot plot shows the upregulated KEGG pathways (p.adjust <0.05) identified using DAVID as enriched in different groups. The size of the dot is based on gene ratio (gene counts) enriched in the pathway, and the color of the dot shows the pathway enrichment significance.

**Figure S11. (A)** List of identified genes in gene ontology terms from RNF20-dependent TFs. **(B)** List of identified genes in gene ontology terms from *cKO*-dependent TFs.
