## Supplementary materials for "Chromatin state transition underlies the temporal changes in gene expression during cardiomyocyte maturation"

**Materials and methods**

**Generation of cardiac-specific and muscle-specific *Rnf20* conditional knockout mice**

Rnf20 conditional knockout targeting vectors were constructed as previously described ^1^ and purchased from EUCOMM/KOMP. Two forward loxP sites were designed and introduced into the construct flanking mouse RNF20 exons 3, 4 and 5, with an upstream neomycin selection cassette (including two Frt sites). The linearized targeting construct was electroporated into C57Bl/6-derived embryonic stem (ES) cells and the clones were screened by Southern blotting and PCR. Four clones out of 389 were positive for the recombination event. The positive clones were injected into C57BL6/J blastocysts. Germ-line transmission was confirmed by coat color as well as PCR analysis of tail DNA. After confirming germ-line transmission of the targeted (*Neo*) allele, the resultant heterozygous *Rnf20^+/flox-Neo^* mice were mated to transgenic mice expressing the FLP recombinase (ACT-FLPe) to induce excision of the *Neo* marker. *Rnf20^flox/flox^* mice were generated by intercrossing *Rnf20^flox/+^* heterozygous mice. Homozygous Rnf 20^flox/flox^ mice were mated to *Mesp1-Cre* ^2^, *Mck-Cre* ^3^ or *Hsa-Cre* ^4^ lines to generate Rnf 20^flox/+^; *Mesp1*-Cre mice, Rnf 20^flox/+^; *Mck*-Cre mice and Rnf 20^flox/+^; *Hsa*-Cre mice. The F1 generations were crossed with homozygous Rnf 20^flox/flox^ mice to generate Rnf 20^flox/flox^; *Mesp1*-Cre mice, Rnf 20^flox/flox^; *Mck*-Cre mice and Rnf 20^flox/flox^; *Hsa*-Cre mice, respectively. The genotypes of the mice were determined by western blot and PCR.

**Animal phenotyping studies**

All animal experiments, including euthanasia protocols, were conducted according to the Guide for the Care and Use of Laboratory Animals, and were approved by the Animal Committee of Institute of Cellular and Organismic Biology, Academia Sinica. Animals were weighed at different time-points between 3 and 10 weeks of age and compared directly to their sex-matched littermates. For prenatal and mortality curve studies, timed breedings were performed, and pregnancy was determined by detection of a vaginal plug (E0.5). The time of birth was closely monitored, and newborns were counted and genotyped within one week after birth. Mice were sacrificed and tissues were harvested for gDNA, RNA and protein extraction at the indicated time-points.

**Cardiomyocyte dissociation**

Ventricular myocytes were obtained by enzymatic dissociation. Mice were anaesthetized by inhalation of isoflurane. Deep anesthesia was confirmed by lack of response to otherwise painful stimuli. The mouse was then euthanized, and the heart was surgically removed via thoracotomy and placed in a Langendorff column. The isolated hearts were perfused sequentially at a constant flow rate of 3 ml/min with Ca^2+^-free solution containing (in mM): 120.4 NaCl, 4.7 KCl, 1.2 MgSO_4_, 0.6 Na_2_HPO_4_, 0.6 KH_2_PO_4_, 4.6 NaHCO_3_, 10 KHCO_3_, 10 HEPES, 30 Taurine and 5.5 glucose, pH 7.0 adjusted with HCl and then an enzyme solution (with collagenase B, D and protease type XIV, Sigma-Aldrich) for 7-9 min. Perfusate temperature was maintained at 37°C. After digestion, ventricles were cut into small pieces, and gently minced by gentle mechanical agitation with a Pasteur pipette. The isolated cardiomyocytes were suspended in 10 ml stop buffer (Ca^2+^-free perfusion buffer with 5% bovine calf serum), and the Ca^2+^ concentration was increased gradually to 1.0 mM. Cardiomyocytes were kept in KB solution containing 85 mM K-glutamate, 30 mM K_2_HPO_4_, 5 mM sodium pyruvate, 0.5 mM EGTA, 20 mM Taurine, 5 mM MgCl_2_, 5 mM Creatine, 2 mM Na_2_ATP, 10 mM BDM and 20 mM Glucose, pH 7.3. Cells were used within 8 h of isolation.

**Histology**

For morphometric analyses, the animals were euthanized at the ages indicated. Hearts were fixed in 10% formaldehyde solution in phosphate buffered saline (PBS), embedded in paraffin, and cut into 10 μm thick sections. Sections were stained with H&E (Hematoxylin and eosin) according to the manufacturer’s instructions. Briefly, paraffin embedded sections were incubated at 60°C for 30 min and rehydrated. Slides were then incubated in Bouin’s solution at 60°C for 1 h, followed by washing and immersion in Weigert’s working hematoxylin solution for 10 min. After repeated washes, slides were immersed in Biebrich Scarlet–Acid Fuchsin solution for 5 min and phosphomolybdic acid solution for 10 min prior to incubation in Aniline blue for 5 min. After a wash, slides were incubated in 1% acetic acid for 1 min, rinsed, dehydrated and mounted with Permount Mounting Medium.

**Electrocardiogram recordings**

Mice were anesthetized with 1.5% isoflurane in 700 ml O_2_ per min via a nose cone (following induction in a chamber containing isoflurane 4–5% in oxygen). Rectal temperature was monitored continuously and maintained at 37–38°C using a heat pad. Three lead ECG (leads I, II, III) were recorded from sterile needle electrodes inserted subcutaneously in each forelimb and hindlimb. The signal was acquired and analyzed using a digital acquisition and analysis system (Power Lab 8/30; AD Instruments, Dunedin, New Zealand). ECG parameters were quantified 1–2 min after anesthesia induction in order to stabilize the trace. Standard three-lead surface ECG recordings were performed continuously for 15 min. The data were digitized and stored for off-line analysis using LabChart Pro software (AD Instruments). QT interval was defined as the time elapsed from the beginning of the major deflection representing the QRS to the end of the secondary slow deflection. QT intervals were corrected for RR interval by the equation (QTc = QT/(RR/100)1/2). Analysis was performed on lead II electrocardiograms.

**Echocardiography**

Two-dimensional echocardiography was performed using a Philips iE33 ultrasound imaging system (Philips Medical Systems, Best, Netherlands) equipped with a 7-15 MHz linear array transducer on P14 to P42 (n = 5 per group) mice. Briefly, after induction of anesthesia in a chamber containing 4-5% isoflurane in oxygen, the mouse was positioned on a heat pad in order to maintain its body temperature at 37–38°C. Anesthesia was maintained with 1.5% isoflurane in 700 ml O_2_ per min via a nose-cone. Heart rate was maintained between 350 and 600 beats per min. After two-dimensional long- and short-axis images of the left ventricular (LV) were obtained, M-mode traces were acquired for measurement of the LV chamber dimensions at the diastole and systole, as well as the wall thickness. Echocardiography-derived LV mass, fractional shortening (FS), and ejection fraction (EF) were recorded. Measurements were averaged from five consecutive cardiac cycles.

**Preparation of tissue extracts and immunoblotting**

Freshly isolated hearts were rinsed with cold PBS and immediately frozen in liquid Nitrogen or homogenized in RIPA buffer (containing 150 mM NaCl, 1% Triton X-100, 0.5% Sodium deoxycholate, 0.1% SDS and 50 mM Tris-HCl (PH 8.0)) with protease inhibitors. Supernatants were collected after centrifugation at 13,200 rpm for 5 min and stored in aliquots at -80°C until use. For western blotting experiments, protein extracts (80 μg of total protein) were separated on 8% or 15% polyacrylamide gels and blotted onto PVDF membranes (Millipore). Membranes were blocked with blocking buffer (5% nonfat dry milk, 10 mM of Tris-HCl, pH 7.6, 150 mM NaCl, and 0.1% Tween 20) and incubated with the primary antibody at 4°C overnight. After incubation with peroxidase-conjugated secondary antibodies, proteins were visualized using enhanced chemiluminescence reagents (Millipore) and detected using an ImageQuant LAS 4000 mini system (GE Healthcare Life Sciences). Densitometric analyses were performed using ImageJ software. The protein levels of β-Actin were used to normalize the results. The primary antibodies included mouse monoclonal antibodies: anti-β-Actin (AC-15, Novus; 1:2,000), anti-GAPDH (GTX100118, GeneTex; 1:10,000), anti-RNF20 (21625-1-AP, Proteintech; 1:2,000) and mitochondrial marker antibody sampler kit (#8674, Cell Signaling).

**Immunofluorescence staining**

For immunofluorescence staining, mice were euthanized and perfused with 4% paraformaldehyde in PBS. The hearts were excised, immediately fixed in 4% paraformaldehyde for 20 min and then equilibrated in 30% sucrose at room temperature. The samples were then embedded into Tissue-Tek OCT compound (Fisher Scientific), and frozen tissues sections were cut (10 μm) and collected. The sections were post-fixed in 2% paraformaldehyde, blocked with 2% bovine serum albumin, and incubated with primary antibodies at 4°C overnight. The primary antibodies included mouse monoclonal antibodies: anti-α-actinin (clone EA-53, Sigma-Aldrich; 1:200.), anti-connexin43 (clone CXN-6, Sigma-Aldrich; 1:200), anti-desmoplakin (clone DP2.15, Millipore Corporation; 1:200), anti-pan cadherin (C3678; Sigma-Aldrich; 1:500), and anti-Vinculin (clone hVIN-1, Sigma-Aldrich; 1:500). After washing in PBS, sections were incubated with secondary antibodies, including Alexa Fluor 488/568 goat anti-mouse or anti-rabbit antibodies (Invitrogen). Counterstaining was performed using 0.5 μg/ml of Hoechst 33342 (Cell Signaling Technology). Fluorescent signals were visualized using a Zeiss LSM700 confocal microscope.

**2D-Transmission electron microscope (TEM)**

Mouse ventricle tissue was diced into small blocks in a fixative mixture of glutaraldehyde (1.5%) and paraformaldehyde (1.5%) in phosphate buffer at pH 7.3. The procedure was performed as described previously ^5^. Ultrathin sections were cut, mounted, post-stained, and observed using a FEI TECNAI G2 F20 S-TWIN electron microscope (Electron Microscope Core Facility, Institute of Cellular and Organismic Biology, Academia Sinica).

**Mitochondria DNA quantification and cristae density measurement**

qPCR analysis for mitochondrial DNA was performed as previously described ^6^. In brief, DNA was purified from freshly frozen heart with Proteinase K digestion and subsequent phenol/chloroform extraction. Mitochondrial DNA (mtDNA) was quantified with SYBR green PCR Master Mix and Roche LightCycler 480 (Roche). Primers used included the following: mtDNA-F: CCCAT TCCAC TTCTG ATTAC C, mtDNA-R: ATGAT AGTAG AGTTG AGTAG CG, nucDNA-F: GTACC CACCT GTCGT CC, nucDNA-R: GTCCA CGAGA CCAAT GACTG. The relative mtDNA copy number was calculated as the ratio of mtDNA copies to nuclear DNA copies. The relative fold-change was then calculated based on the ΔΔ*C*t method. Cristae density was also measured as previously reported ^6^.

**Mitochondria enzymatic activity assay and ATP measurement**

Samples were prepared as previously described ^7^. Briefly, hearts (50 mg) were minced and homogenized in 1.5 ml of homogenization buffer (10 mM MOPS, 1.0 mM EDTA, 210 mM mannitol, and 70 mM sucrose, pH 7.4) using a Polytron homogenizer (low setting, 3 s). The homogenate was centrifuged at 500*g* for 5 min (4°C), and the supernatant was filtered through cheese cloth. Protein concentration of the lysate was measured. Samples were diluted to 1,000 μg/ml in 10 mM Tris-Hcl (pH 7.0) containing 250 mM sucrose and stored at -80°C.

For NADH oxidase activity, 10 μl heart homogenate was used. NADH-supported electron transport was assessed spectrophotometrically as the rate of NADH consumption (340 nm, *ε* = 6,200 M^−1^cm^−1^) following the addition of NADH (Working solution: 50 mM phosphate buffer, 145 μM NADH and 0.5 mM EDTA). Cytochrome C oxidase activity was measured according to the instructions included in the colorimetric assay kit (#K287-100, BioVision). ATP content was measured using the ENLITEN® ATP Assay System (FF2000, Promega).

**RNA-seq and TO-GCN analysis**

Total RNA was isolated from hearts of P2 to P28 mice with Trizol, according to the manufacturer’s instructions. RNA was further column purified with a RNeasy MinElute kit (#28004, Qiagen) prior to sequencing. The isolated RNA was then prepared for sequencing using the Ovation SoLo or Ovation Universal RNA-seq System kits (Tecan), according to the manufacturer's instructions. Libraries were sequenced on the Illumina NextSeq high output platform at 400 million reads with single reads length 75 in Institute of Molecular Biology of Academia Sinica. Approximately 20 million reads per sample were generated. Sequencing results were demultiplexed and converted to FASTQ format using Illumina Bcl2FastQ software. Quality Control (QC) for the RNA-Seq reads was assessed using FastQC software.

After quality filtering according to the Illumina pipeline, paired-end reads were aligned to the mm10 reference genome with mouse gene annotation (Ensembl build 97) using HISAT2 ^8^. Gene expression levels in FPKM (fragments per kilobase of exon per million fragments mapped) were quantified using StringTie ^9^. For subsequent analyses, nuclear protein-coding genes with FPKM equal to or greater than one in at least one sample were selected. For gene expression level comparisons between samples, upper quantile normalization was applied. Differentially expressed genes (DEGs) between samples were called using NOISeq ^10^ with probability > 0.99. The annotation of mouse TF was downloaded from animalTFDB 3.0 [https://doi.org/10.1093/nar/gky822]. The TO-GCNs (time-ordered gene coexpression networks) were constructed using the TO-GCN pipeline (https://github.com/petitmingchang/TO-GCN) ^11^. In this study, the input data were expression of 986 TF genes from a time series of transcriptomes derived from heart tissue of control (WT) and *Rnf20* mutants (*cKO*). The Pearson correlation coefficient (PCC) values for all TF-TF pairs were calculated for WT and *cKO* separately, and cutoffs of positive co-expression PCC ≥ 0.91 (p-value < 0.05) for both conditions were determined. WT+ and *cKO*+ TO-GCNs were constructed using Plagl1 and Hey2 as the initial TF genes, respectively.

**ATAC-seq and TFs footprinting analysis**

ATAC-seq on frozen mouse heart tissues was performed according to a modified ATAC-seq (Omni-ATAC) protocol from a previous study ^12^. ATAC-seq raw data was processed using the published ATAC-seq pipeline ATAC-seq ^13^. Adaptor-trimmed and quality-filtered reads were mapped to the mm10 reference genome using Bowtie2 ^14^ with options “--very-sensitive -X 2000”. PCR duplicates, reads aligned to mitochondrion genome, and non-unique aligned reads were removed. MACS2 ^15^ was applied for peak calling. Annotation and pathway enrichment analyses for peak associated genes were conducted using ChIPseeker ^16^. TF motif finding and footprinting analyses were conducted using HINT-ATAC ^17^ and TOBIAS ^18^, respectively.

**Statistical methods**

Data were compiled and are shown as mean ± standard error of the mean (SEM). Data were evaluated using unpaired, two-tailed *t* tests (95% confidence interval) or two-way analyses of variance with post hoc analysis using GraphPad Prism4 software (GraphPad Inc, San Diego, CA). *P-*values <0.05 were considered significant.

**Data and software availability**

The accession number for the RNA-seq and ATAC-seq reported in this paper is BioProject ID: PRJNA745479.

(<https://dataview.ncbi.nlm.nih.gov/object/PRJNA745479?reviewer=2mbh9mjvqasvas1dquls89091r>)

**[References]**

1.	Skarnes, W.C. *et al.* A conditional knockout resource for the genome-wide study of mouse gene function. *Nature* **474**, 337-42 (2011).

2.	Saga, Y. *et al.* MesP1: a novel basic helix-loop-helix protein expressed in the nascent mesodermal cells during mouse gastrulation. *Development* **122**, 2769-78 (1996).

3.	Johnson, J.E., Wold, B.J. & Hauschka, S.D. Muscle Creatine-Kinase Sequence Elements Regulating Skeletal and Cardiac-Muscle Expression in Transgenic Mice. *Molecular and Cellular Biology* **9**, 3393-3399 (1989).

4.	Miniou, P. *et al.* Gene targeting restricted to mouse striated muscle lineage. *Nucleic Acids Res* **27**, e27 (1999).

5.	Wu, C.Y. *et al.* A persistent level of Cisd2 extends healthy lifespan and delays aging in mice. *Hum Mol Genet* **21**, 3956-68 (2012).

6.	Puente, B.N. *et al.* The oxygen-rich postnatal environment induces cardiomyocyte cell-cycle arrest through DNA damage response. *Cell* **157**, 565-79 (2014).

7.	Nakada, Y. *et al.* Hypoxia induces heart regeneration in adult mice. *Nature* **541**, 222-227 (2017).

8.	Kim, D., Paggi, J.M., Park, C., Bennett, C. & Salzberg, S.L. Graph-based genome alignment and genotyping with HISAT2 and HISAT-genotype. *Nat Biotechnol* **37**, 907-915 (2019).

9.	Pertea, M. *et al.* StringTie enables improved reconstruction of a transcriptome from RNA-seq reads. *Nat Biotechnol* **33**, 290-5 (2015).

10.	Tarazona, S. *et al.* Data quality aware analysis of differential expression in RNA-seq with NOISeq R/Bioc package. *Nucleic Acids Res* **43**, e140 (2015).

11.	Chang, Y.M. *et al.* Comparative transcriptomics method to infer gene coexpression networks and its applications to maize and rice leaf transcriptomes. *Proceedings of the National Academy of Sciences of the United States of America* **116**, 3091-3099 (2019).

12.	Corces, M.R. *et al.* An improved ATAC-seq protocol reduces background and enables interrogation of frozen tissues. *Nat Methods* **14**, 959-962 (2017).

13.	Zuo, Z. *et al.* ATAC-pipe: general analysis of genome-wide chromatin accessibility. *Brief Bioinform* **20**, 1934-1943 (2019).

14.	Langmead, B. & Salzberg, S.L. Fast gapped-read alignment with Bowtie 2. *Nat Methods* **9**, 357-9 (2012).

15.	Zhang, Y. *et al.* Model-based analysis of ChIP-Seq (MACS). *Genome Biol* **9**, R137 (2008).

16.	Yu, G., Wang, L.G. & He, Q.Y. ChIPseeker: an R/Bioconductor package for ChIP peak annotation, comparison and visualization. *Bioinformatics* **31**, 2382-3 (2015).

17.	Li, Z. *et al.* Identification of transcription factor binding sites using ATAC-seq. *Genome Biol* **20**, 45 (2019).

18.	Bentsen, M. *et al.* ATAC-seq footprinting unravels kinetics of transcription factor binding during zygotic genome activation. *Nat Commun* **11**, 4267 (2020).
